# Taxonomically informed scoring enhances confidence in natural products annotation

**DOI:** 10.1101/702308

**Authors:** Adriano Rutz, Miwa Dounoue-Kubo, Simon Ollivier, Jonathan Bisson, Mohsen Bagheri, Tongchai Saesong, Samad Nejad Ebrahimi, Kornkanok Ingkaninan, Jean-Luc Wolfender, Pierre-Marie Allard

## Abstract

Mass spectrometry (MS) hyphenated to liquid chromatography (LC)-MS offers unrivalled sensitivity for metabolite profiling of complex biological matrices encountered in natural products (NP) research. With advanced platforms LC, MS/MS spectra are acquired in an untargeted manner on most detected features. This generates massive and complex sets of spectral data that provide valuable structural information on most analytes. To interpret such datasets, computational methods are mandatory. To this extent, computerized annotation of metabolites links spectral data to candidate structures. When profiling complex extracts spectra are often organized in clusters by similarity via Molecular Networking (MN). A spectral matching score is usually established between the acquired data and experimental or theoretical spectral databases (DB). The process leads to various candidate structures for each MS features. At this stage, obtaining high annotation confidence level remains a challenge notably due to the high chemodiversity of specialized metabolomes.

The integration of additional information in a meta-score is a way to capture complementary experimental attributes and improve the annotation process. Here we show that integrating unambiguous taxonomic position of analyzed samples and candidate structures enhances confidence in metabolite annotation. A script is proposed to automatically input such information at various granularity levels (species, genus, and family) and weight the score obtained between experimental spectral data and output of available computational metabolite annotation tools (ISDB-DNP, MS-Finder, Sirius). In all cases, the consideration of the taxonomic distance allowed an efficient re-ranking of the candidate structures leading to a systematic enhancement of the recall and precision rates of the tools (1.5 to 7-fold increase in the F1 score). Our results clearly demonstrate the importance of considering taxonomic information in the process of specialized metabolites’ annotation. This requires to access structural data systematically documented with biological origin, both for new and previously reported NPs. In this respect, the establishment of an open structural DB of specialized metabolites and their associated metadata (particularly biological sources) is timely and critical for the NP research community.

## 1 Introduction

Specialized metabolites define the chemical signature of a living organism. Sessile organisms such as plants, sponges and corals, but also microorganisms (bacteria and fungi), are known to biosynthesize a wealth of such chemicals. These molecules can play a role as defense or communication agents (Brunetti et al., 2018). However, the functional role of these metabolites for the producer is not always fully understood. The comprehension of these functions is one of the goals of chemical ecology and recent research illustrates the importance of specialized metabolism (Hoffmann et al., 2018; Schmitt et al., 1995). Throughout history, humans have been relying on plant derived products for a variety of purposes: housing, feeding, clothing and, especially, medication. Our therapeutic arsenal is in fact deeply dependent on the chemistry of natural products (NPs) whether they are used in mixtures, purified forms or for hemi-synthetic drug development. After a period of disregard by the pharmaceutical industry, NPs are the object of a renewed interest (Shen, 2015). Today, technological and methodological advances allow to establish more precise views of the chemo-biologic aspects of life. In the field of NPs, advances in genomics and metagenomics now allow the anticipation of the chemical output directly from environmental DNA (Alanjary et al., 2019; Craig et al., 2009). Developments in metabolite profiling by mass spectrometry (MS) grant access to large volumes of high-quality spectral data from minimal amount of samples and appropriate data analysis workflows allow to efficiently mine such data (Wolfender et al., 2019). Initiatives such as the Global Natural Products Social (GNPS) molecular networking (MN) (Wang et al., 2016) project offer both a living MS repository and the possibility to establish MN organizing MS data. Despite such advancements, **metabolite identification** remains a major challenge for both NP research and metabolomics (Kind et al., 2018). Metabolite identification of a novel compound requires physical isolation of the analyte followed by complete NMR acquisition and three-dimensional structural establishment via X-ray diffraction or chiroptical techniques. For previously described compounds, metabolite identification implies complete matching of physicochemical properties between the analyte and a standard compound (including chiroptical properties). Metabolite identification is thus a tedious and labor-intensive process, which should ideally be reserved to novel metabolites’ description. Any less complete process should be defined as **metabolite annotation**. By definition, metabolite annotation can be applied at a higher throughput and offers an effective proxy for the chemical characterization of complex matrices. This process includes dereplication (the annotation of previously described molecules prior to any physical isolation process) and allows focusing isolation and metabolite identification efforts on potentially novel compounds only (Newman, 2017).

Given its sensitivity, selectivity and structural determination potential, MS is a tool of choice for metabolite annotation in complex mixtures. High resolution mass spectrometers (HRMS) affording high mass accuracy (ppm range) are now routinely used (Eliuk and Makarov, 2015; Lommen et al., 2011; Olsen et al., 2005). As a result of this evolution, unambiguous molecular formula (MF) can be assigned in most cases. However, the isomeric nature of many NPs indicates that MF determination is a necessary but non-sufficient step in the metabolite annotation process. Acquisition of fragmentation spectra (tandem MS (MS/MS) or MSn) offers a way to gain structural insights on analytes. Such analytical approach (physically breaking down the analyte in constitutive building blocks to infer its general structure) is analog to DNA (genomics) or protein sequencing (proteomics). However, compared to those two fields, a notable difference lies within the non-polymeric nature of small metabolites, which partly explains the difficulties encountered in linking a fragmentation spectrum to a chemical structure in metabolomics. The dependence of the fragmentation behavior of analytes on the mass spectrometer geometry adds further complexity to the interpretation of MS data (Johnson and Carlson, 2015). To interpret such data, a common approach consists in comparing experimental spectra to established spectral libraries. In the specific case of GC-MS, the reproducibility of the electron ionization (EI) mediated gas-phase fragmentation process allowed to establish huge spectral libraries (e.g. NIST) comparable across laboratories worldwide. The situation is quite different for soft ionization electrospray (ESI)-MS spectra acquired in LC-MS. The lack of standardization, fragmentation variability and difficult access to standards has complicated the establishment of robust experimental libraries. As a result, it is estimated that no more than 25,000 unique compounds are present in such libraries (Dührkop et al., 2015). Various computational MS solutions have appeared to overcome such limitations and link experimental spectra to chemical structures. They can be classified into experimental rule-based strategies (MassHunter, Agilent Technologies), combinatorial fragmentation strategies (MetFrag, (Ruttkies et al., 2016)), machine learning based approaches using stochastic Markov modelling (CFM-ID, (Allen et al., 2014; Djoumbou-Feunang et al., 2019)) or predicting fragmentation trees (Sirius) (Böcker et al., 2009; Dührkop et al., 2019). Computationally demanding ab initio calculations, modeling the gas-phase fragmentation process, have also been proposed (Bauer and Grimme, 2016). The output of such tools is, in general, a list of candidate molecules ranked according to a score. Such score can be based on a single measure (e.g. spectral similarity in CFM-annotate) (Allen et al., 2014) or integrate combined parameters (MS-Finder, Sirius) (Dührkop et al., 2019; Tsugawa et al., 2016, 2019).

In our view, increased confidence in specialized metabolite annotation can be achieved by the establishment of a meta-scoring system capturing the similarity of diverse attributes shared by the queried analytes and candidate structures (Allard et al., 2017). Such meta-score could for example consider 1) spectral similarity 2) taxonomic distance between the producer of the candidate compound and the annotated biological matrix 3) structural consistency within a cluster and 4) physico-chemical consistency. See Fig 1 for a conceptual overview of such metascore. The rationale behind comprehensive scoring systems is that orthogonal information (not directly related to spectral comparison) should strengthen the metabolite annotation process. This has been illustrated by using additional information, such as the number of literature references related to a candidate structure and basic retention time scoring based on logP in MetFrag 2.2 (Ruttkies et al., 2016). Recently, retention order prediction has been integrated to an MS/MS prediction tool and also provided increased performance in metabolite annotation (Bach et al., 2018). Another example is the Network Annotation Propagation (NAP) approach, which takes advantage of the topology of a MN to proceed to a re-ranking of annotated candidates within a cluster where structural consistency is expected (da Silva et al., 2018).

**Figure 1.**
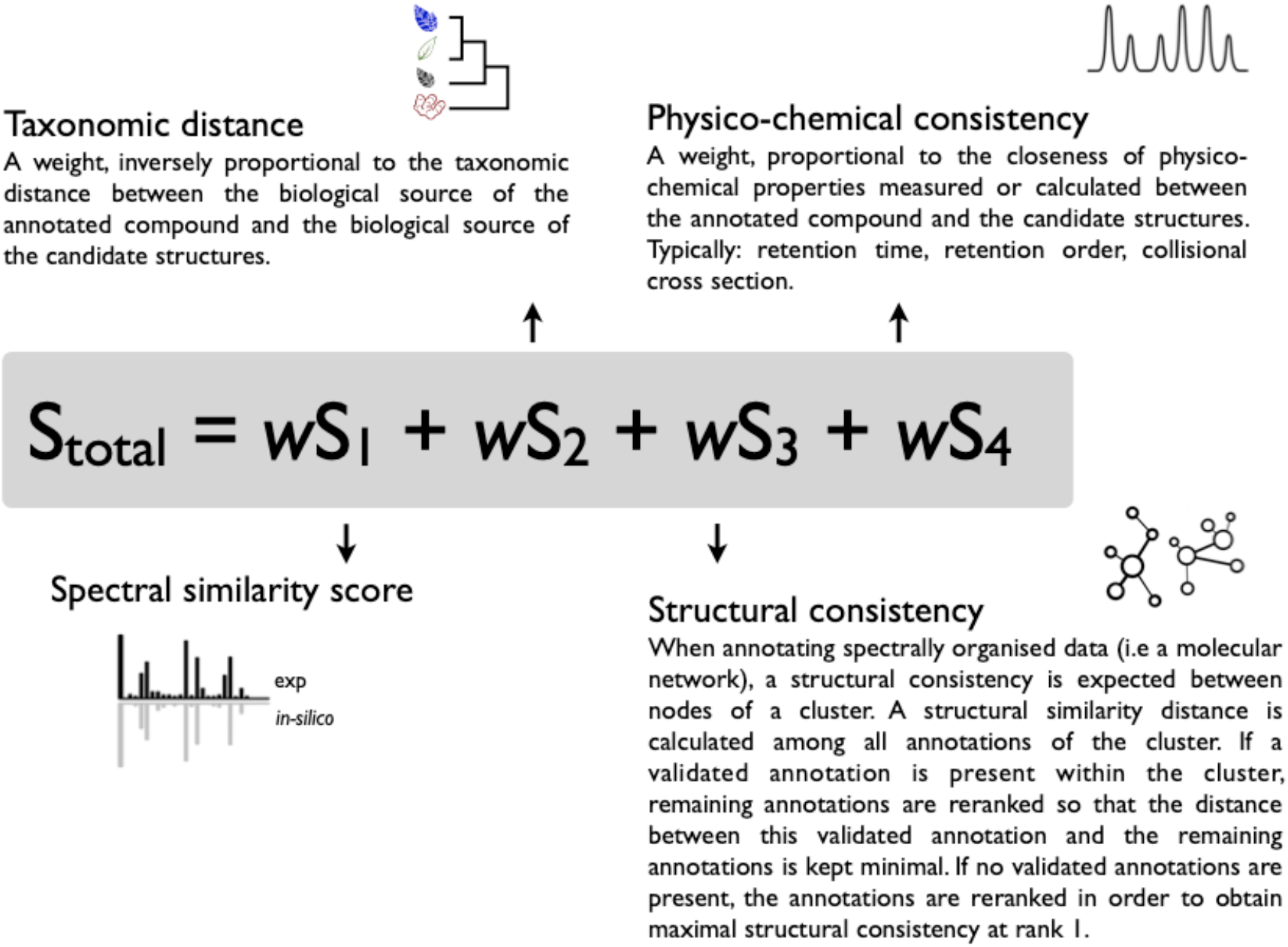
Conceptual overview of a possible meta-scoring system for specialized metabolite annotation incorporating 1) spectral similarity 2) taxonomic distance between the biological source of the queried spectra and candidate annotations in the database (DB) 3) structural consistency within a cluster (see **(da Silva et al., 2018)**) and 4) physico-chemical consistency (see **(Ruttkies et al., 2016)** and **(Bach et al., 2018)**). A factor (w) should allow to attribute relative weight to individual scores and modulate their contribution to the overall score.

To the best of our knowledge, the automated inclusion of the taxonomic dimension has not been considered in current metabolite annotation strategies. The central hypothesis of this work is directly inferred from the characteristics of the specialized metabolome. Unlike primary metabolites, which are mostly ubiquitous compounds central to cell functioning, specialized metabolites are, by definition, strongly linked to the taxonomic position of the producing organisms. Such principles were already formulated in 1816 by de Candolle (de Candolle, 1828) who postulated that 1) *Plant taxonomy would be the most useful guide to man in his search for new industrial and medicinal plants* and 2) *Chemical characteristics of plants will be most valuable to plant taxonomy in the future*. More than 200 years later both principles have been largely verified, as recently illustrated (Hoffmann et al., 2018; Kang et al., 2019).

It thus appears desirable to consider taxonomic information when describing the chemistry of an organism. A taxonomic *filtering* process could be used to limit a DB to compounds previously isolated in organisms situated within a given taxonomic distance from the biological source of the analyte to annotate. However, results of chemotaxonomic studies also highlight the presence of broadly distributed metabolites. For example, liriodenine (<u>MUMCCPUVOAUBAN-UHFFFAOYSA-N</u>) is a widely distributed alkaloid produced by more than 50 distinct biological sources, it is found in over 30 genus belonging to 13 botanical families. Convergent biosynthetic pathways offer intriguing example of unrelated species, shaped by evolution, that end up producing similar classes of compounds (Pichersky and Lewinsohn, 2011). To adjust for the annotation of such compounds, a more tolerant *weighted scoring system* allowing, both, to consider spectral similarity and taxonomic information while conserving the independence of the individual resulting scores appears as a better solution.

In the frame of this study we propose such taxonomically informed scoring system and benchmark the impact of taxonomic distance consideration on a set of 2107 identified molecules using three different computational mass spectrometry metabolite annotation tools (ISDB-DNP, MS-Finder and Sirius).

## 2 Results

### 2.1 Conception of the taxonomically informed scoring system

The constituents of specialized metabolomes, as expression products of the genome, should reflect the taxonomical position of the producing organism. Our initial working hypothesis is thus that *the attribution of a weight, inversely proportional to the taxonomic distance between the biological source of the queried analyte and the one of the candidate structures, should be a valuable input into the metabolite annotation process.*

The proposal of the taxonomically informed scoring system is thus to complement the initial score given to candidate structures by existing metabolite annotation tool by such a weight. To this end, the initial score should first be normalized. Then, weights, inversely proportional to the taxa level difference (family < genus < species) are given. Such weights are attributed when an exact match is observed between biological source denominations at the following taxa levels: family, genus and species. The weight corresponding to the shortest taxonomic distance is then added to the initial score. Candidates are further re-ranked according to the newly weighted score. The general outline of the taxonomically informed scoring system is presented in Fig. 2. In this study, no phylogenetic distances within taxa (e.g. family, genus or species) were considered.

**Figure 2.**
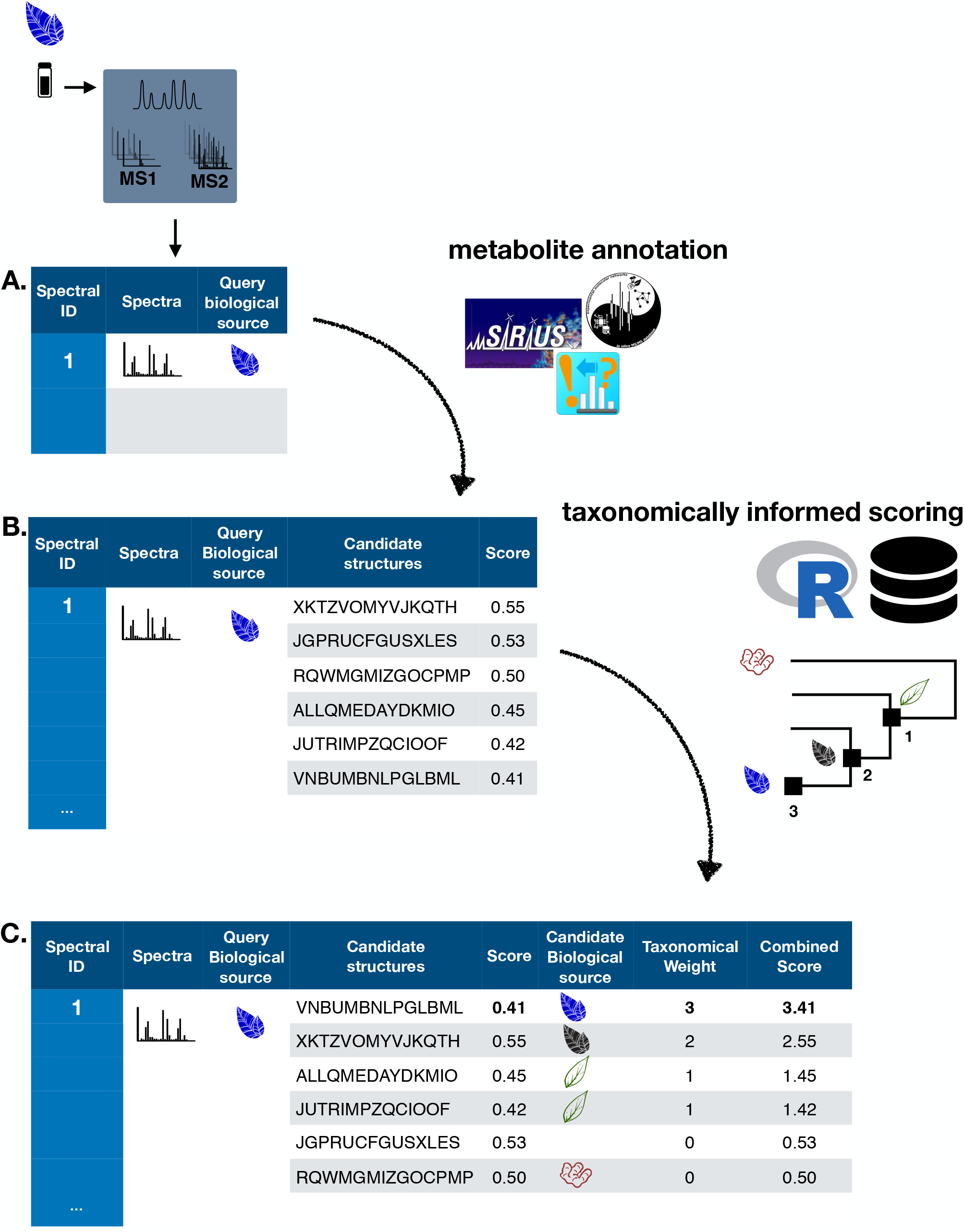
General outline of the taxonomically informed scoring system. Candidates structures are complemented with their biological sources at the family, genus and species level, when available. A weight, inversely proportional to the taxonomic distance between the biological source of the standard compound and the one of the candidate compounds is given when the biological source of the candidate structures matches the biological source of the standard at the family, genus and species level, respectively. The maximal weight for each candidate is then added to its spectral score to yield a weighted spectral score. Finally, candidates are re-ranked according to the weighted spectral score.

In order to apply the taxonomically informed scoring system in a generic manner, the initial scores given by the metabolite annotation tools were rescaled to obtain values ranging from 0 (worst candidate) to 1 (best candidate). The weights, given according to the taxonomic distance between the biological source of the queried spectra and the one of the candidate compounds, were integrated in the final score by a sum. This choice allows to keep independence between individual components of the metascore (see Fig. 1). Since the boundaries of the candidates’ normalized score in a given dataset are defined (0 to 1), the minimal weight to be applied to the worst candidate for it to be ranked at the first position after weighting is 1. Following our initial hypothesis, a weight of 1 was thus given if a match between biological sources was found at the family taxa level. In the case were the initial maximal score (1) would be given to a candidate and added to a weight corresponding to a match at the family level (1), a weight of at least 2 should be given for a candidate having the worst score to be ranked above. A weight of 2 was thus given if a match between biological sources was found at the genus taxa level. Following the same logic, a weight of 3 was given for matches between biological sources at the species level.

### 2.2 Benchmarking the influence of taxonomic information consideration in metabolite annotation

#### 2.2.1 Establishment of a benchmarking dataset

In order to establish the importance of considering taxonomic information during the metabolite annotation we needed to construct a dataset constituted by molecular structures, their MS/MS fragmentation spectra acquired under various experimental conditions and their biological sources in the form of a fully resolved taxonomical hierarchy. This dataset, denominated hereafter “benchmarking dataset”, was built by combining a curated structural/biological sources dataset (obtained from the DNP) and a curated structural/spectral dataset (obtained from GNPS libraries) as described in Material and Methods. Below are the results of each step.

##### 2.2.1.1 Structural and biological sources dataset

The prerequisite to apply a taxonomically informed scoring in a metabolite annotation process is to dispose of the biological source information of *1)* the queried spectra and *2)* the candidate structures. To the best of our knowledge, there is currently no freely available database (DB) compiling NP structures and their biological sources down to the species level. For this study, we thus exploited the DNP that is commercially available and allows export of structures and biological sources as associated metadata. For example, the output for the biological source field of pulsaquinone, “Constit. of the roots of *Pulsatilla koreana*.”, is Plantae | Tracheophyta | Magnoliopsida | Ranunculales | Ranunculaceae | *Pulsatilla* | *Pulsatilla cernua*. Within the Global Names index, biological sources resolved against the Catalogue of Life source were kept resulting in 219,800 entries with accepted scientific names and full, homogeneous, taxonomy up to the kingdom level. See Material and Methods for example and details concerning the curation process.

##### 2.2.1.2 Structural and spectral dataset

The GNPS libraries agglomerate a wide and publicly available ensemble of MS/MS spectra coming from various analytical platforms and thus having different levels of quality (Wang et al., 2016). These spectral libraries were used as representative source of diverse experimental MS/MS spectra to evaluate the annotation improvement that could be obtained by applying taxonomically informed scoring system. For this purpose, all GNPS libraries and publicly accessible third-party libraries were retrieved online (https://gnps.ucsd.edu/ProteoSAFe/libraries.jsp) and concatenated as a single spectral file containing 66646 individual entries. The pretreatment described in corresponding Material and Methods section, yielded a dataset of 40138 structures with their experimental associated MS/MS acquired on different platforms.

##### 2.2.1.3 Structural, spectral and biological sources dataset (benchmarking set)

To implement the taxonomically informed metabolite annotation process, it is required that both *1)* the queried spectra and *2)* the candidate structures biological sources are equally resolved (i.e. using the accepted denomination) at the taxonomic level. It was thus necessary to build a spectral dataset for which each entry had a unique structure and a properly documented biological source, which constituted the benchmarking set. For this, the structural and spectral datasets was matched against the structural and biological source dataset, following the procedure detailed in Material and Methods. The full processing resulted in a dataset of 2107 individual entries (characterized NPs with no stereoisomers distinction and a unique biological source associated), which was used for the rest of this study.

#### 2.2.2 Evaluation of the improvement of metabolite annotation on the benchmarking set

In order to assess the importance of considering taxonomic information in the annotation process, the output of three different computational mass spectrometry-based metabolite annotation solutions were submitted to the taxonomically informed scoring. The 2107 spectra of the benchmarking set were queried using these three tools according to parameters detailed in the Material and Methods section. This resulted in three different outputs constituted by a list of candidates for each entry of the benchmarking set.

##### 2.2.2.1 Metabolite annotation tools used

- ***ISDB-DNP*** The first tool, denominated hereafter ISDB-DNP (In Silico DataBase – Dictionary of Natural Products) is metabolite annotation strategy that we previously developed (Allard et al., 2016). This approach is focused on specialized metabolites annotation and is constituted by a pre-fragmented theoretical spectral DB version of the DNP. The in silico fragmentation was performed by CFM-ID, a software using a probabilistic generative model for the fragmentation process, and a machine learning approach for learning parameters for this model from MS/MS data (Allen et al., 2015). CFM is, to our knowledge, the only computational solution currently available able to generate a theoretical spectrum with prediction of fragment intensity. The matching phase between experimental spectra and the theoretical DB is based on a simple spectral similarity measure (cosine score) performed using Tremolo as a spectral library search tool (Wang and Bandeira, 2013). The scores are reported from 0 (worst candidate) to 1 (best candidate).
- ***MS-Finder*** The second tool is MS-Finder. This in silico fragmentation approach considers multiple parameters such as bond dissociation energies, mass accuracies, fragment linkages and various hydrogen rearrangement rules at the candidate ranking phase (Tsugawa et al., 2016). The resulting scoring system range from 1 (worst candidate) to 10 (best candidate).
- ***Sirius*** The third tool to be used is Sirius 4.0. It is considered as a state-of-the-art metabolite annotation solution, which combines molecular formula calculation and the prediction of a molecular fingerprint of a query from its fragmentation tree and spectrum (Dührkop et al., 2019). Sirius uses a DB of 73,444,774 unique structures for its annotations. The resulting score is a probabilistic measure ranging between negative infinity and 0 (best candidate).

##### 2.2.2.2 Computation of the taxonomically informed scoring system

R scripts were written to perform *1)* cleaning and standardization of the outputs, *2)* taxonomically informed scoring and re-ranking and *3)* summary of the resulting annotations. First, the outputs were standardized to a table containing on each row: the unique spectral identifier (CCCMSLIB N°) of the queried spectra, the short InChIKey of the candidate structures, the score of the candidates (within the scoring system of the used metabolite annotation tool), the biological source of the standard compound and the biological source of the candidate structures. As described in section 2.1, a weight, inversely proportional to the taxonomic distance between the biological source of the annotated compound and the biological source of the candidate structure, was given when both matched at the family (weight of 1), genus (weight of 2) or species level(s) (weight of 3). A sum of this weight (1-3) and the original score (0 to 1) yielded the taxonomically weighted score. This taxonomically weighted score was then used to re-rank the candidates.

###### Taxonomically unweighted results

Using each tools’ initial scoring system, on the 2107 experimental MS/MS spectra constituting the benchmarking set, the ISDB-DNP returned 214 (10.2 %) correct annotations at rank 1, Sirius 975 (46.3 %) and MS-Finder 180 (8.5 %). The total number of unique correct annotations ranked first covered by ISDB-DNP, Sirius and MS-Finder prior to taxonomically informed scoring reached 1110 or 52.7 % of the benchmarked library. Out of these, 29 (less than 1.4 %), were common to all three tools, indicating the interest of considering various annotation tools when proceeding to metabolite annotation. See Venn diagram in **Fig. 3** illustrating the complementarity of the returned annotations. Within the list of all candidates, the ISDB-DNP returned 1750 correct annotations, Sirius 1589 and MS-Finder 574. The ROC curves (Supplementary Material S4) outline the number of correct hits outside first rank and indicate remaining improvement potential.

**Figure 3.**
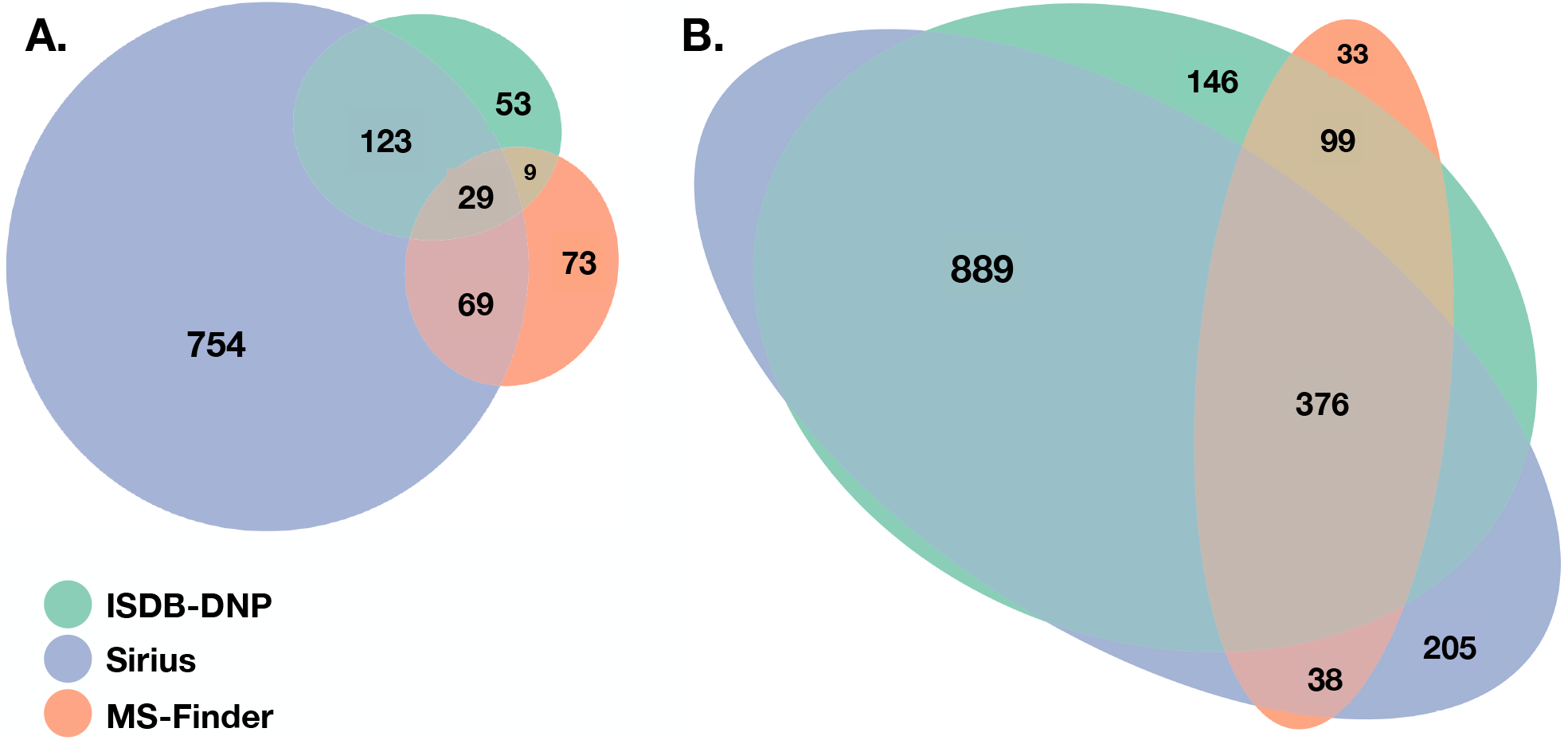
Venn diagrams representing common and unique correct annotations of each tool at rank 1 before (A.) and after (B.) taxonomically informed scoring.

###### Taxonomically weighted results

After weighting and reranking with the taxonomically informed score, the number of correct annotations at rank 1 increased to 1510, 1508 and 546, respectively for ISDB-DNP, Sirius and MS-Finder. The annotation complementarity between the tools is highlighted in Fig. 4. The total number of correct annotations covered by all ISDB-DNP, Sirius and MS-Finder after taxonomically informed scoring reached 1786 or 84.8 % of the benchmarked library. Interestingly, a more than 10-fold increase after taxonomically informed scoring was also observed for the correctly annotated metabolite commonly returned by the three tools 376 (17 %). It is to be noted that no stereoisomer distinction could be performed since all correct matches were assessed by short InChIKey comparison.

**Figure 4.**
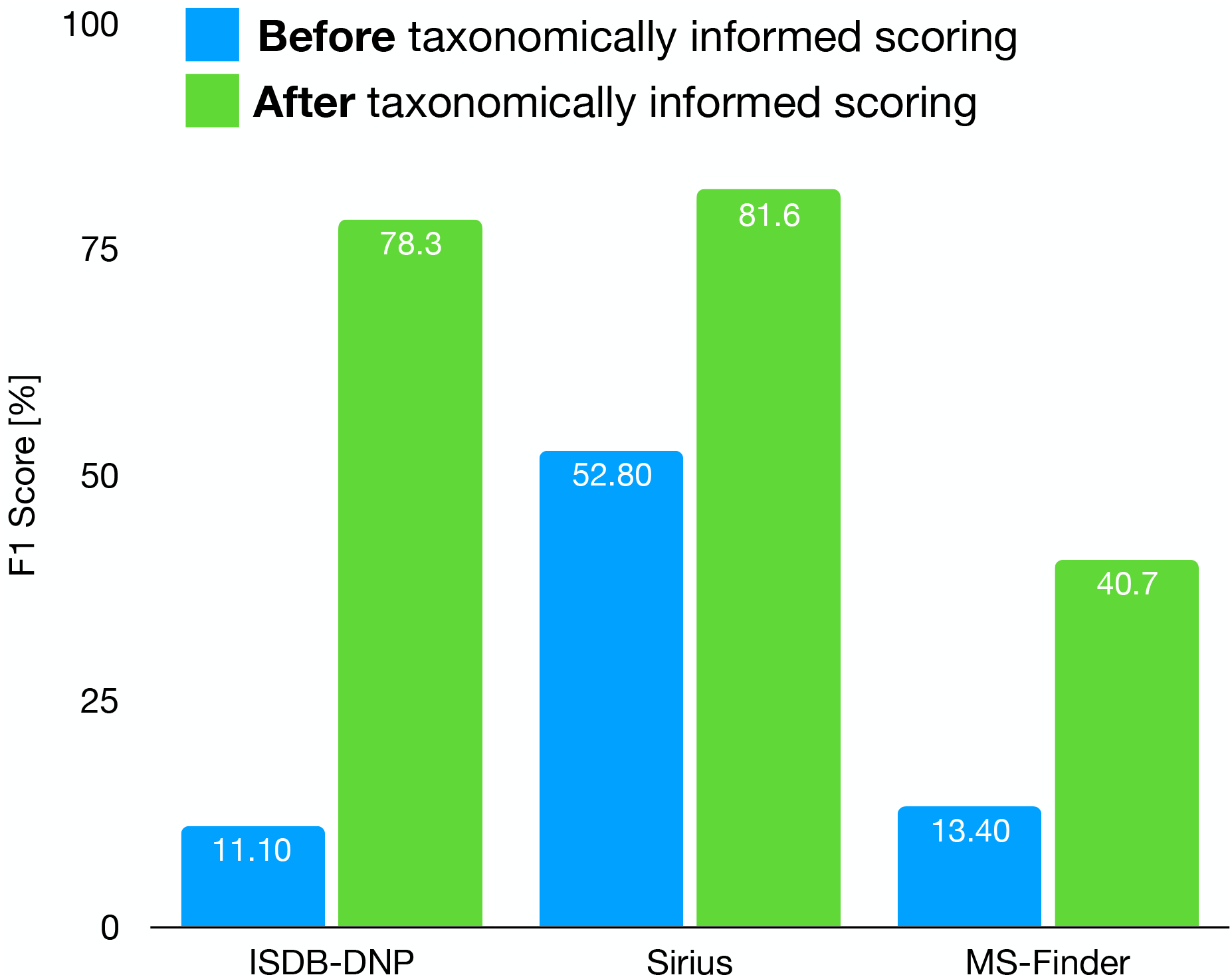
Influence of the taxonomically informed scoring on the F1 score of each metabolite annotation tool.

In order to evaluate the impact of the taxonomically informed scoring system, the F1 score was used. More details can be found in Material and Methods. The results of this treatment on the outputs of the three metabolite annotation tools are displayed on **Fig. 4**. The taxonomically informed scoring stage led to a systematic increase of the F1 score for the benchmarked tools. This increase was of 7 fold (ISDB-DNP), 1.5 (Sirius) and 3 fold (MS-Finder).

### 2.3 Optimization of weights combination for the taxonomically informed scoring

In order to verify our initial hypothesis and define the optimal weights combination (at the family, genus and species taxa level) to be applied for taxonomically informed scoring we proceeded to a global optimization of the taxonomically informed scoring function.

To this end, the taxonomic information related to candidate’s annotation was artificially degraded. This step allowed to mimic a “real life” case in which correct candidate annotation’s taxonomic metadata are not necessarily complete down to the species level. Using the procedure detailed in the corresponding Material and Methods section, the algorithm was applied four times on the four randomized datasets. It quickly converged (100 iterations) towards a global maximum (max 1126 hits, see Fig 5.). The global optimal parameters were found to be 0.81, 1.62 and 2.55 for the family, genus and species taxa level, respectively. Such scores are dependent on the nature and completeness of the employed taxonomic metadata. However, the results obtained when applying the Bayesian optimization on the annotation sets (for which taxonomic metadata was randomly degraded) indicated that optimal results were systematically obtained when *the attributed weights were inversely proportional to the taxa hierarchical position*, thus confirming our initial hypothesis.

**Figure 5.**
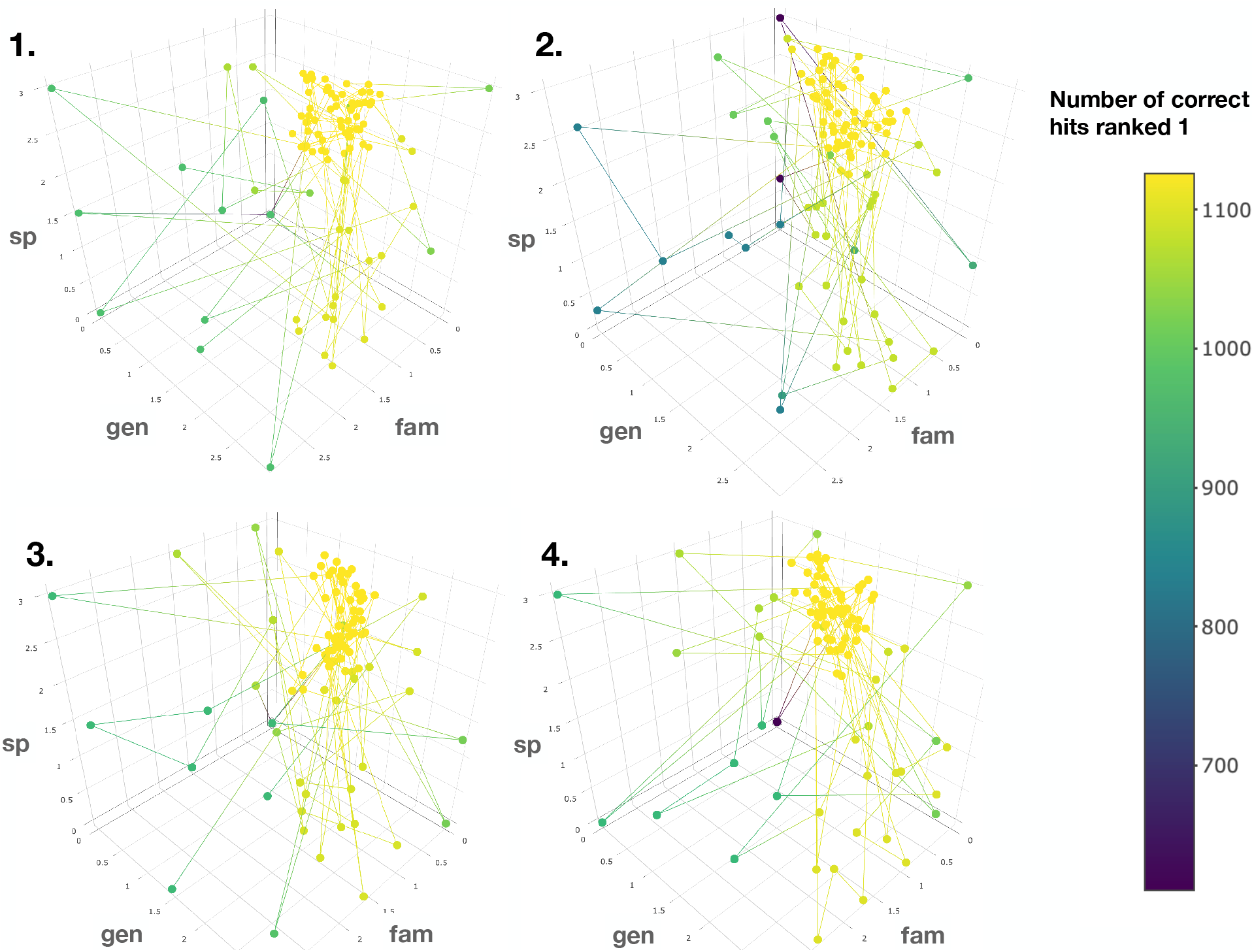
Results of the Bayesian optimization converge toward the optimal weights combination required for a maximal number of correct annotation ranked at the first position. This is observed for four randomly degraded training sets (first optimization round displayed). The results confirm that the applied weights should be *inversely proportional to the taxonomic distance between the biological source associated to the queried spectra and the biological source of the candidate structures*.

### 2.4 Application of the taxonomically informed scoring to the annotation of metabolites from *Glaucium* sp

The impact of the taxonomically informed scoring was evaluated in a real case example for the annotation of metabolites from *Glaucium* species (Papaveraceae family). Three species, *G. grandiflorum*, *G. fimberilligerum* and *G. corniculatum* were studied. The ethyl acetate and methanolic extracts of the three species were profiled by UHPLC-HRMS in positive ionization mode using a data-dependent MS2 acquisition. After appropriate data treatment and molecular networking (see corresponding Material and Methods section), the taxonomically informed scoring was used to proceed to the re-ranking of the candidates according to the respective taxonomic position of their biological sources and return the top 5 hits. We especially focused on the two major compounds (MS signal intensity) of *G. grandiflorum.* These were feature *m/z* 342.1670 at 1.42 min and *m/z* 356.1860 at 1.83 min. A weight of 0.81, 1.62 and 2.55 was thus given to candidates for which the biological source was found to be Papaveraceae at the family level, *Glaucium* at the genus level and *G. grandiflorum* at the species level, and respectively. The results of the taxonomically informed scoring annotation for feature *m/z* 342.1670 at 1.42 min are presented in Table 1. (see SI for annotation results concerning feature *m/z* 356.1860 at 1.83 min). Both features were targeted within the extract and, after isolation, the structure of their corresponding compound was determined by 1D and 2D NMR measurements (see spectra in Supplementary Material S1 and S2). The NMR spectra matched the literature reported spectra for predicentrine (Guinaudeau et al., 1979) in case of feature *m/z* 342.1670 at 1.42 min and glaucine (Huang et al., 2004) for feature *m/z* 356.1860 at 1.83 min. In both cases, the candidate structure proposed via the taxonomically informed scoring annotation at rank 1 was found to be correct. With the classical spectral matching process, the correct candidates were initially ranked at position 9 and 7 for predicentrine and glaucine, respectively (see Table 1 and Supplementary Material S5 and S6).

**Table 1.**
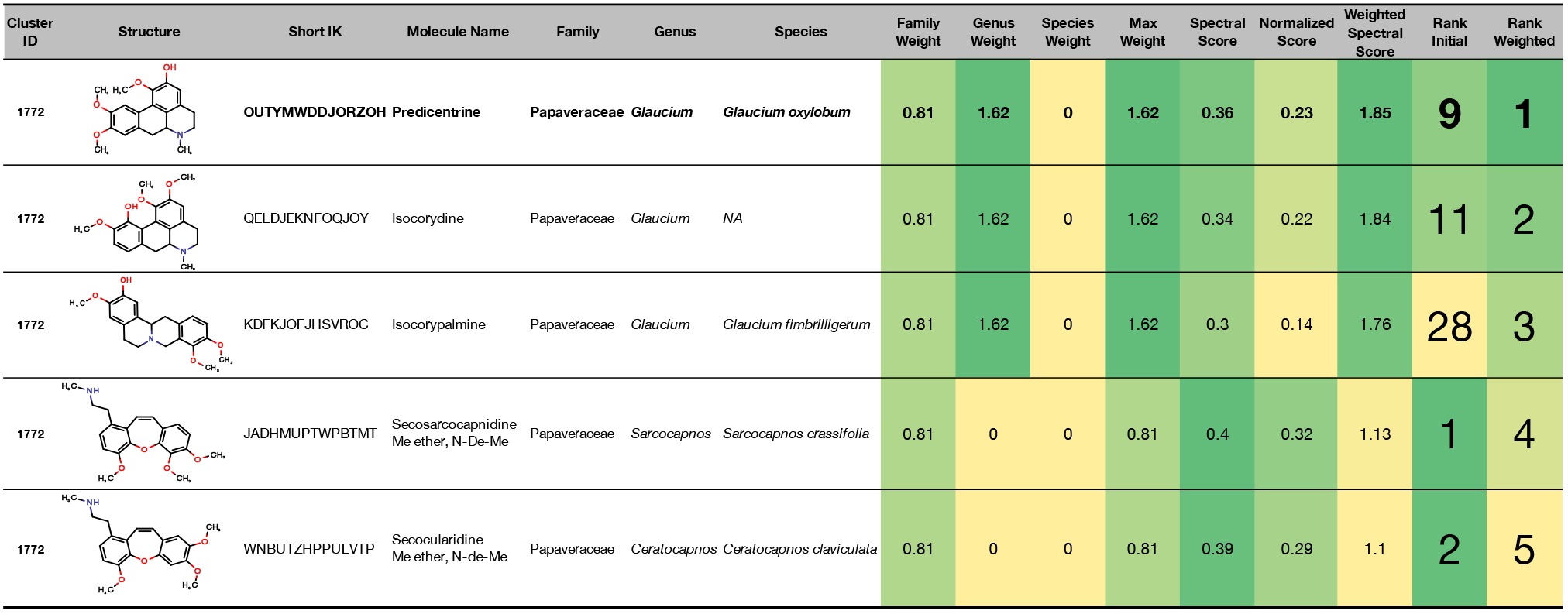
Output of the taxonomically informed scoring annotation using ISDB-DNP for feature *m/z* 342.1670 at 1.42 min. Predicentrine, which is the correct annotation and was initially ranked at the 9th position, is now ranked at the first position.

## 3 Discussion

Given the complexity of natural extracts, the isolation of specialized metabolites thereof is a long and tedious process which should ideally be carried on NPs of high value only (e.g. novel or bioactive compounds). On the other hand, metabolite annotation in complex extracts is also required for metabolomics studies, compositional assessment of phyto-preparation or chemotaxonomical studies. The annotation process is thus key for many aspects of NP research. This process can be boiled down to the comparison of attributes (e.g. exact mass, MF, fragmentation spectra) of the queried analyte to attributes of candidate structures present in a DB. When HRMS and appropriate heuristic filters are used, the establishment of the molecular formula (MF) of the analyte is relatively straightforward (Kind and Fiehn, 2007). However, this is not sufficient to proceed to metabolite annotation given the isomeric nature of numerous NP. Over all the compounds reported in the DNP, less than 10 % (9.46 %) have a unique chemical formula. For example, 147 occurrences correspond to the chemical formula C_16_H_14_O_4_. With a MS1 analysis relying on exact mass only, no ranking between those isomers is possible. By filtering the biological sources, the number of possible structures can be significantly reduced, but can still lead to multiple isomers. In the case of *Bauhinia manca* (synonym of *Bauhinia guianensis*), there are 5 entries corresponding to this formula. Taking both MS/MS fragmentation and taxonomic position into account, the confidence into the annotation process is notably enhanced leading in this case to the correct annotation at rank 1 (see Supplementary Material S3). Comparison of spectra to experimental or in silico fragmented DBs, allows to attribute a score to candidate structures and, thus, to discriminate isomeric molecules. However, MF and fragmentation spectra are not the only attributes which can be compared in the metabolite annotation process. Specialized metabolites, as products of biosynthetic clusters themselves part of the genome, are tightly linked to the taxonomic position of the producing organisms (Ernst et al., 2019b; Hoffmann et al., 2018). Here, we demonstrate that the biological source and, more precisely, the taxonomic distance between the biological source of the queried compound and the biological source of the candidate structures is also a valuable attribute to integrate into the metabolite annotation process. We show that such information can be considered in a taxonomically informed scoring system and automatically applied to the outputs of different computational metabolite annotation programs. The consideration of taxonomic information was shown to systematically improve the F1 score of the evaluated solutions (ISDB-DNP, Sirius, MS-Finder) with a 1.5 to 7-fold increase. The advantage of considering such information in the metabolite annotation process are thus observed independently of the tools and their associated structural DBs. This benchmarking was carried to evaluate the importance of considering taxonomical information during the metabolite annotation process. It is not meant to compare the performances of the tools. Indeed, all compounds of the benchmarking set were present in the DNP, and the ISDB-DNP tool, which is by definition backed by the same database is thus favored. Furthermore filters (for the selection of [M+H]+ adducts and for the filtering MS/MS spectra for the 500 most intense peaks) were applied to meet restriction of the ISDB-DNP and MS-Finder, respectively. Finally, a number of entries (197) of the benchmarking dataset were found to have large mass difference (> 0.01 Da) between their experimental parent ion mass and their calculated exact mass. For example, cevadine [M+H]+ (<u>CCMSLIB00004689734</u>) had an experimental parent ion mass of 632.386 Da, while its calculated exact mass is 591.3407 Da (C_32_H_49_NO_9_). Of course, such erroneous entries cannot be identified by the computational metabolite annotation tools. The list of these problematic entries is available online (<u>Problematic_entries.csv</u>). It is also important to keep in mind that such metabolite annotation tools are incapable of discriminating stereoisomers.

In the two cases studied in this work (annotation of standard compounds and annotation of analytes originating from a single biological sources), the attribution of the biological source to the queried features is straightforward. However, a more complicated configuration can appear: the annotation of aligned MS data coming from an extract library. In this particular case a feature can belong to multiple biological sources. It is thus necessary to link features to multiple extracts (biological sources) for example via MS signal intensity.

The results of the optimization on datasets for which taxonomical information had been randomly degraded at multiple taxa level indicated, for the ISDB-DNP results, that the optimal combination of weights was 0.81, 1.62 and 2.55 (family, genus and species taxa level, respectively). Such results should however be taken with caution, and not as absolute optimal values, as such optimization process are heavily dependent on the learning sets. The optimization however indicates that the best results were achieved when the assigned weights were inversely proportional to the taxonomic distance between the biological sources of, both, the queried spectra and the candidate structures.

It is important to note that such taxonomically informed scoring system will mostly benefit the annotation process of specialized metabolites and not ubiquitous molecules (e.g. coming from the primary metabolism) for obvious reasons. Furthermore, it heavily depends on the availability and quality of DBs compiling structures and their biological sources reported as a fully and homogeneously resolved taxonomy. To the best of our knowledge, such DBs are not publicly available at the moment. The NPAtlas (https://www.npatlas.org/) is an interesting initiative of the Linington lab, however biological sources information down to the species level is only accessible in query mode and the DB is limited to 20440 metabolites of microbial origin only. The Dictionary of Natural Products (DNP), which we used in this study is the widest compilation of structure/biological sources pairs, but is only available commercially. Furthermore, the biological sources are reported as a free text field (codes are available only for the family taxa levels and above), thus requiring tedious standardization and name resolving. It is therefore important for the community to start the systematic reporting of biological sources, together with spectral and structural information, when documenting novel metabolites. In fact, the reporting of newly described biological occurrence should be encouraged even for previously described metabolites. However, the policy of most journals in NP research is to accept for publication only description of novel and bioactive structures, which hinders these potentially informative reports. The GNPS spectral libraries (https://gnps.ucsd.edu/ProteoSAFe/libraries.jsp) and MassIVE repositories (https://massive.ucsd.edu/ProteoSAFe/static/massive.jsp) appear as the optimal place, at the moment, to compile and share NP spectral and structural information. But, if free text comments can complement the documentation of an entry, no standardized fields are available to report the biological sources of the uploaded spectra. The creation of such a feature, ideally directly linking the entered biological sources to existing taxonomy backbones such as GBIF (https://www.gbif.org) or Catalogue of Life (http://www.catalogueoflife.org), would be extremely useful. A recent initiative, the Pharmacognosy Ontology (PHO), that builds on the 50 years of development of NAPRALERT (https://www.napralert.org) is aimed at providing a Free and Open resource that will link taxonomical, chemical and biological data (http://ceur-ws.org/Vol-1747/IP12_ICBO2016.pdf).

The taxonomically informed scoring presented here only takes into account the identity between the biological sources, at different taxa level, of the query compounds and the ones of the candidates structures. Taking into account a more precise phylogenetic position within or across taxa, for example via the calculation of taxonomic distinctiveness indexes (Clarke and Warwick, 1998; Weikard et al., 2006), could offer a more accurate distance and eventually improve such taxonomically informed metabolite annotation process. Of course, and in addition to correct and systematic biological sources occurrence reporting in dedicated DBs, it is of utmost importance to count on the expert knowledge of trained taxonomists specialized in the classification of biodiversity. But it appears that today, sadly, these people are missing (Ajmal Ali and Choudhary, 2011; Drew, 2011)

As proposed in the introduction (see also Fig. 1), an ideal metabolite annotation process should consider a maximal number of attributes when matching the queried analyte to candidate structures present in a DB. It is clear that MS1 based annotation is not sufficient when annotating NP given their isomeric nature, and MS/MS based workflow are known the standard way to proceed to their annotation. We have shown here that complementing the MS/MS comparison with taxonomical information is beneficial for the metabolite annotation process. Others groups have demonstrated the advantages of integrating information such as retention order (Bach et al., 2018), the topology of a molecular network (relatedness within a cluster) in order to rerank candidates proposed by MS/MS based annotation (da Silva et al. 2018). Efforts remains to be done towards the establishment of a global meta-score (see. Fig 1) and problematics such as the individual weights to attribute to each individual component of a metascore will likely appear, but integrating the maximal number of metadata available when proceeding to metabolite annotation should only be beneficial for such process. Overall, considering the relations across attributes by contextualization approaches (molecular networking, biosynthetic coherence, chemotaxonomic relationships) should be particularly efficient to annotate the chemistry of living systems (Allard et al., 2018). Various approaches have been proposed to exploit structural (or biosynthetic) relationships among metabolites and further organize the producing extracts (Junker, 2018; Liu et al., 2017) and interesting developments will appear once robust metabolite annotation solution are coupled to comprehensive DBs compiling structures and their biological sources. Indeed, specialized metabolome annotation could be a novel way to infer the taxonomic position of an unknown sample, just as valid as a genetic sequencing. This could offer new possibilities for taxonomists and application in the field of phyto-pharmaceutical quality control for the identification of adulterations in a similar way to MALDI-TOF based approaches now identifying bacterial strains in mixtures (Yang et al., 2018).

## 4 Conclusion

Efficient characterization of specialized metabolomes is a key challenge in metabolomics and NP chemistry. Now, technical advances allow to access large amounts of informative data, requiring ad hoc computational solutions for their interpretation. The metabolite annotation process, can be resumed to the comparison of attributes of the queried features against attributes of the candidate structures. This process can benefit from the information complementary to the classically used MS fragmentation fingerprints. Ideally, the quantification of multiples attributes similarity (or dissimilarity) should be integrated within a metascoring system (see Fig 1). Here, we demonstrate that the consideration of the taxonomic distance separating the biological sources of both, the queried analytes, and the candidate structures, can drastically improve the efficiency (recall and precision rates) of existing computational metabolite annotation solutions. Metabolite annotation is crucial to guide chemical ecology research or drug discovery projects. We therefore go in the direction of the first of De Candolle’s assumptions (*“Plant taxonomy would be the most useful guide to man in his search for new industrial and medicinal plants”*). His correlated postulate (*“Chemical characteristics of plants will be most valuable to plant taxonomy in the future”*) has been validated and exciting research in this direction is being carried at the moment (MSXXX). Metabolite annotation can benefit from taxonomy and taxonomical relationships can be inferred from precise metabolite characterization. Efforts in both directions should thus fuel a virtuous cycle of research aiming to better understand Life and its chemistry.

These progresses will however depend on the establishment of open data repositories systematically compiling structures, spectra and associated biological sources but also capturing organization of these objects at various levels (structural and spectral similarity, taxonomical relationships). The Pharmacognosy Ontology project should provide a valuable framework to document and contextualize such valuable data (http://ceur-ws.org/Vol-1747/IP12_ICBO2016.pdf).

## 5 Material and Methods

### 5.1 Outline and implementation of the taxonomically informed scoring system

To evaluate the importance of considering taxonomic information in the annotation process, three different computational mass spectrometry-based metabolite annotation tools were used (namely, ISDB-DNP, MS-Finder and Sirius). This resulted in three different outputs constituted by a list of candidates returned by each tool for the entries of the benchmarking dataset. These candidates were ranked according to the scoring system of each tool. R scripts in the form of markdown notebooks were written to perform *1)* cleaning and standardization of the outputs (<u>TaxoCleaneR.Rmd</u>) *2)* taxonomically informed scoring and re-ranking (<u>TaxoWeighter.Rmd</u>) and *3)* analysis of the results (<u>TaxoDesigner.Rmd</u>). First, the outputs were standardized to a table containing on each row: a unique spectral identifier (CCMSLIB N°) of the queried spectra, the short InChIKey of the candidate structures, the score of the candidates (within the scoring system of the used metabolite annotation tool), the biological source of the standard compound and the biological source of the candidate structures. As described in section 2.1, a weight, inversely proportional to the taxonomic distance between the biological source of the annotated compound and the one biological source of the candidate structure, was given when an exact match was found between both biological sources at the family, genus or/and species level(s). A sum of this weight and the original score yielded the taxonomically weighted score. This taxonomically weighted score was then used to re-rank the candidates. See Fig. 2 for a schematic overview of the taxonomically informed scoring process.

### 5.2 Dataset preparation

- **Structural and biological sources dataset** In the Dictionary of Natural Products (v 27.1), taxonomic information appears in two fields. The Biological Source field, which is constituted by a free text field reporting occurrence of a specific compound and the Compound Type field which reports various codes corresponding to molecule classes or taxonomic position at the family level. As an example, for the entry corresponding to larictrin 3-glucoside (ODXINVOINFDDDD-UHFFFAOYSA-N), the Biological Source field indicates “Isol. from *Larix* spp., *Cedrus* sp. and other plant spp. Constit. of *Vitis vinifera* cv. Petit Verdot grapes and *Abies amabilis.*” and the Compound Type field indicates “V.K.52600 W.I.40000 W.I.35000 Z.N.50000 Z.Q.71600” suggesting that biological sources are found in the Phyllocladaceae (Z.N.50000) and Vitaceae family (Z.Q.71600). The biological source information is reported in a non-homogeneous way and multiple biological sources are reported in the same row. In order to extract taxonomic information out of the free text contents, we used the *gnfinder* program (https://github.com/gnames/gnfinder). Gnfinder takes UTF8-encoded text as inputs and returns back JSON-formatted output that contains detected scientific names. It automatically detects the language of the text and uses complementary heuristic and natural language processing algorithms to detect patterns corresponding to scientific binomial or uninomial denomination. We used gnfinder forcing for English language detection. In addition to scientific denomination extraction, gnfinder allows to match the detected names against the Global Names index services (https://index.globalnames.org). The preferred taxonomy backbone was set to be Catalogue of Life. This last step allowed to return the full taxonomy down to the entered taxa level. It also allows to resolve synonymy. Since gnfinder is designed to mine free texts, the JSON formatted output indicates the position of the detected name in the original input by character position. A python script was written to output a .csv file with the found name and taxonomy in front of the corresponding input. When multiple biological sources were found for an entry, this one was duplicated in order to obtain a unique structure/biological source pair per row. The script is available online (<u>gnfinder_field_scrapper.py</u>).
- **Structural and spectral dataset** All GNPS libraries and publicly accessible third-party libraries were retrieved online (https://gnps.ucsd.edu/ProteoSAFe/libraries.jsp) and concatenated as a single spectral file (<u>Full_GNPS_lib.mgf</u>) in the .mgf format. A python Jupyter notebook (<u>MGF_lib_filterer.ipynb</u>) was created to filter .mgf spectral file according to specific parameters: maximum and minimum number of fragments per spectrum and defined spectral ID (e.g. CCMLIB N°). All parameters are optional. The spectral file was filtered to retain only entries having at least 6 fragments. For spectra containing more than 500 fragments, only the 500 most intense were kept. A second python Jupyter notebook (<u>GNPS_lib_parser_cleaner.ipynb</u>) was written to proceed to *1)* extraction of relevant metadata (parent ion mass, Smiles, InChI, library origin, source instrument, molecule name and individual spectrum id value (CCMSLIB N°) *2)* filtering entries having at least one structural information associated (Smiles and/or InChI) and corresponding to protonated adducts and *3)* converting structures to their InChIKey, a 27-character hashed version of the full InChI. The InChIKey conversion was realized using the RDKit 2019.03.1 framework (RDKit: Open-source cheminformatics; http://www.rdkit.org). This resulted in a structural dataset (<u>GNPS_lib_structural.tsv</u>) of 40138 entries. The library was further filtered to keep entries which parent masses were comprised between 100 and 1500 Da. Duplicate structures and stereoisomers were removed by keeping distinct InChIKey according to the first layer (first 14 characters) of the hash code. This spectral dataset encompasses spectra acquired on a variety of MS platforms. The codes, input and output data are available on OSF at the following address (https://osf.io/bvs6x/).
- **Structural, spectral and benchmarking dataset (benchmarking dataset)** Once the structural and biological sources dataset and the structural and spectral datasets were prepared (as described above), both were joined in order to attribute a biological source to each spectrum. The scripts used to proceed to the merging step are part of the python Jupyter notebook (<u>GNPS_lib_parser_cleaner.ipynb</u>). Since in most cases it is not expected to differentiate stereoisomers based on their MS spectra, the combination of both datasets was made using the short InChIKey (first 14 characters of the InChIKey) as a common key. In this merging process only, entries having biological source information resolved against the Catalogue of Life and complete down to the species level were retained. However, this merging implies that for a given biological source the information on the 3D aspects of the structure is lost. While this is not an issue for the benchmarking objective of this work the resulting dataset doesn’t constitute a reliable occurrence dataset for annotation that needs stereoisomers to be differentiated. The resulting dataset containing structural, spectral and biological sources information was constituted by 2107 distinct entries. This constituted the benchmarking dataset. The scripts allowing to generate the benchmarking dataset, the associated spectral data (<u>Benchmark_dataset_spectral.mgf</u>), and associated metadata (<u>Benchmark_dataset_metadata.tsv</u>) are available at the following address (https://osf.io/bvs6x/).

### 5.3 Computational metabolite annotation tools

- **ISDB-DNP** The ISDB-DNP (In Silico DataBase – Dictionary of Natural Products) is an approach that we previously developed (Allard et al., 2016). A version using the freely available Universal Natural Products Database (ISDB-UNPD) is available online (http://oolonek.github.io/ISDB/). This approach is focused on specialized metabolites annotation and is constituted by a pre-fragmented theoretical spectral DB version of the DNP. The in silico fragmentation was performed using CFM-ID, a software using a probabilistic generative model for the fragmentation process, and a machine learning approach for learning parameters for this model from MS/MS data (Allen et al., 2015). CFM, is, to the best of our knowledge, the only solution available at the moment allowing to output a spectrum with fragment intensity prediction. The matching phase between experimental spectra and the theoretical DB is based on a plain spectral similarity computation performed using Tremolo as a spectral library search tool (Wang and Bandeira, 2013). The parameters used to proceed to the benchmarking dataset analysis were the following: parent mass tolerance 0.05 Da, minimum cosine score 0.1, no limits for the returned candidates numbers.
- **MS-Finder** This in silico fragmentation approach considers multiple parameters such as bond dissociation energies, mass accuracies, fragment linkages and various hydrogen rearrangement rules at the candidate ranking phase (Tsugawa et al., 2016). The resulting scoring system range from 1 to 10. The parameters used to proceed to the benchmarking dataset analysis were the following: mass tolerance setting: 0.1 Da (MS1), 0.1 Da (MS2); relative abundance cut off: 5 % formula finder settings: LEWIS and SENIOR check (yes), isotopic ratio tolerance: 20 %, element probability check (yes), element selection (O, N, P, S, Cl, Br). Structure Finder setting: tree depth: 2, maximum reported number: 100, data sources (all except MINEs DBs. Total number of structures, 321617.) MS-Finder v. 3.22 was used, it is available at the following address: http://prime.psc.riken.jp/Metabolomics_Software/MS-FINDER/
- **Sirius** Sirius 4.0.1 is considered as a state-of-the-art metabolite annotation solution, which combines molecular formula calculation and the prediction of a molecular fingerprint of a query compound from its fragmentation tree and spectrum (Dührkop et al., 2019). Sirius uses a DB of 73,444,774 unique structures for its annotations. The parameters used to proceed to the benchmarking dataset analysis were the following for Sirius molecular formula calculation: possible ionization [M+H]^+^, instrument: Q-TOF, ppm tolerance 50 ppm, Top molecular formula candidates: 3, filter :formulas from biological DBs. For the CSI:FingerID step, the parameters were the following: possible adducts: [M+H]^+^, filter: compounds present in biological DB, maximal number of returned candidates: unlimited. Sirius 4.0.1 is available at the following address: https://bio.informatik.uni-jena.de/software/sirius/

### 5.4 Results analysis

The F1 score was calculated for each evaluated metabolite annotation tool before and after the weighting step. The F1 score is the harmonic mean of the recall (True Positive / (True Positive + False Negative)) and precision rate (True Positive / (True Positive + False Positive)) of a tool. The True Positive (TP) corresponds to the number of correct candidate annotations at rank 1, the False Positive (FP) to the number of wrong candidate annotations at rank 1, and the False Negative (FN) to the number of correct annotations at rank > 1. The F1 score is then calculated as follows:

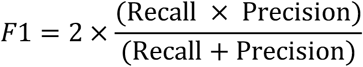

R notebook to analyze the results of the taxonomically informed scoring process and plot the Venn diagrams presented in Fig. 4 are available online (<u>TaxoDesigner.Rmd</u>).

### 5.5 Optimization of the weight combination for the taxonomically informed scoring

In order to establish the optimal weights to be applied for each of the taxonomic distances (family, genus and species), the information related to candidates’ annotations was artificially degraded. For this, the annotation set returned by the ISDB-DNP approach against the benchmarking dataset was randomized. The randomized annotation set was then split into four equal blocks. For the first three blocks, the biological source information was deleted, respectively, at the species level; at the genus and species level; and, finally, at the family, genus and species levels. The fourth block was not modified. Finally, the four blocks were merged back to a unique dataset. The process was repeated four times yielding four datasets with randomly incomplete biological sources. The taxonomic distance informed scoring process was compiled to a unique function taking three arguments (weights given when a match was found at the family, genus and species level, respectively) and outputting the number of correct hits ranked at the first position. A parallelizable Bayesian optimization algorithm (https://github.com/AnotherSamWilson/ParBayesianOptimization) was then used, being particularly suited for the optimization of black box functions for which no formal representation is available (<u>arXiv:1807.02811</u>). The bounds were set between 0 and 3 for the exploration of the three parameters of the function. Number of initial points was set to 10 and the number of iterations to 100. Parameter kappa (κ) was set to 5.152, to force the algorithm to explore unknown areas. The chosen acquisition function was set to Expected Improvement (ei). Epsilon parameter (ε, eps) was set to 0. The whole procedure was run 4 times on the 4 randomized datasets. Best set of parameters were then averaged across the 16 results set. All codes required for this optimization step are available online (<u>TaxoOptimizeR.Rmd</u>).

### 5.6 Chemical analysis and isolation of compounds from *Glaucium* extract

#### 5.6.1 Plant material

The aerial flowering parts of three *Glaucium* species were collected in May and June of 2015 from the northern part of Iran including Mazandaran and Tehran provinces. The samples were identified by Dr. Ali Sonboli, Medicinal Plants and Drugs Research Institute, Shahid Beheshti University, Tehran, Iran. Voucher specimens (MPH-2351 for *G. grandiflorum*, MPH-2352 for *G. fimberilligerum* and MPH-2353 for *G. corniculatum*) have been deposited at the Herbarium of Medicinal Plants and Drugs Research Institute (HMPDRI), Shahid Beheshti University, Tehran, Iran.

#### 5.6.2 Mass Spectrometry analysis

Chromatographic separation was performed on a Waters Acquity UPLC system interfaced to a Q-Exactive Focus mass spectrometer (Thermo Scientific, Bremen, Germany), using a heated electrospray ionization (HESI-II) source. Thermo Scientific Xcalibur 3.1 software was used for instrument control. The LC conditions were as follows: column, Waters BEH C18 50 × 2.1 mm, 1.7 μm; mobile phase, (A) water with 0.1 % formic acid; (B) acetonitrile with 0.1 % formic acid; flow rate, 600 μL.min^−1^; injection volume, 6 μL; gradient, linear gradient of 5−100 % B over 7 min and isocratic at 100 % B for 1 min. The optimized HESI-II parameters were as follows: source voltage, 3.5 kV (pos); sheath gas flow rate (N_2_), 55 units; auxiliary gas flow rate, 15 units; spare gas flow rate, 3.0; capillary temperature, 350.00 °C, S-Lens RF Level, 45. The mass analyzer was calibrated using a mixture of caffeine, methionine − arginine − phenylalanine − alanine − acetate (MRFA), sodium dodecyl sulfate, sodium taurocholate, and Ultramark 1621 in an acetonitrile/methanol/water solution containing 1% formic acid by direct injection. The data-dependent MS/MS events were performed on the three most intense ions detected in full scan MS (Top3 experiment). The MS/MS isolation window width was 1 Da, and the stepped normalized collision energy (NCE) was set to 15, 30 and 45 units. In data-dependent MS/MS experiments, full scans were acquired at a resolution of 35 000 FWHM (at *m/z* 200) and MS/MS scans at 17 500 FWHM both with an automatically determined maximum injection time. After being acquired in a MS/MS scan, parent ions were placed in a dynamic exclusion list for 2.0 s.

#### 5.6.3 MS data pretreatment

The MS data were converted from the .RAW (Thermo) standard data format to .mzXML format using the MSConvert software, part of the ProteoWizard package (Chambers et al., 2012). The converted files were treated using the MzMine software suite v. 2.38 (Pluskal et al., 2010).

The parameters were adjusted as following: the centroid mass detector was used for mass detection with the noise level set to 1.0E6 for MS level set to 1, and to 0 for MS level set to 2. The ADAP chromatogram builder was used and set to a minimum group size of scans of 5, minimum group intensity threshold of 1.0E5, minimum highest intensity of 1.0E5 and *m/z* tolerance of 8.0 ppm. For chromatogram deconvolution, the algorithm used was the wavelets (ADAP). The intensity window S/N was used as S/N estimator with a signal to noise ratio set at 25, a minimum feature height at 10000, a coefficient area threshold at 100, a peak duration ranges from 0.02 to 0.9 min and the RT wavelet range from 0.02 to 0.05 min. Isotopes were detected using the isotopes peaks grouper with a m/z tolerance of 5.0 ppm, a RT tolerance of 0.02min (absolute), the maximum charge set at 2 and the representative isotope used was the most intense. An adduct (Na^+^, K^+^, NH_4_^+^, CH_3_CN^+^, CH_3_OH^+^, C_3_H_8_O^+^ (IPA^+^)) search was performed with the RT tolerance set at 0.1 min and the maximum relative peak height at 500%. A complex search was also performed using [M+H]^+^ for ESI positive mode, with the RT tolerance set at 0.1 min and the maximum relative peak height at 500%. Peak alignment was performed using the join aligner method (*m/z* tolerance at 8 ppm), absolute RT tolerance 0.065 min, weight for *m/z* at 10 and weight for RT at 10. The peak list was gap-filled with the same RT and *m/z* range gap filler (*m/z* tolerance at 8 ppm). Eventually the resulting aligned peaklist was filtered using the peak-list rows filter option in order to keep only features associated with MS2 scans.

#### 5.6.4 Molecular networks generation

In order to keep the retention time, the exact mass information and to allow for the separation of isomers, a feature-based MN (https://bix-lab.ucsd.edu/display/Public/Feature+Based+Molecular+Networking) was created using the .mgf file resulting from the MzMine pretreatment step detailed above. Spectral data was uploaded on the GNPS molecular networking platform. A network was then created where edges were filtered to have a cosine score above 0.7 and more than 6 matched peaks. Further edges between two nodes were kept in the network if and only if each of the nodes appeared in each other’s respective top 10 most similar nodes. The spectra in the network were then searched against GNPS' spectral libraries. All matches kept between network spectra and library spectra were required to have a score above 0.7 and at least 6 matched peaks. The output was visualized using Cytoscape 3.6 software (Shannon et al., 2003). The GNPS job parameters and resulting data are available at the following address (https://gnps.ucsd.edu/ProteoSAFe/status.jsp?task=a475a78d9ae8484b904bcad7a16abd1f).

#### 5.6.5 Taxonomically informed metabolite annotation

The spectral file (.mgf) and attributes metadata (.clustersummary) obtained after the MN step were annotated using the DNP-ISDB with the following parameters: parent mass tolerance 0.005 Da, minimum cosine score 0.2, maximal number of returned candidates: 50. An R script was written to proceed to the taxonomically informed scoring on GNPS outputs and return an attribute table which can be directly loaded in Cytoscape. The script is available here (<u>Taxo_WeighteR_UseR.Rmd</u>).

#### 5.6.6 Isolation of predicentrine and glaucine from *G. grandiflorum*

The air-dried, ground and powdered plant materials (500 g) was successively extracted by solvents of increasing polarities (hexane, ethyl acetate and methanol), 4 × 5.0 L of each solvent (48 h). An aliquot of each ethyl acetate and methanolic extract was submitted to C18 SPE (eluted with 100% MeOH), dried under nitrogen flow and redissolved at 5 mg/ml in MeOH for LC-MS analysis. The methanolic extract of *G. grandiflorum* was concentrated under reduced pressure, then dried with a nitrogen flow until complete evaporation of the residual solvent yielding 50 g of extract. An aliquot (5 g) was subjected to a VLC in order to eliminate sugars and other very polar compounds. A 250 mL sintered-glass Buchner funnel connected to a vacuum line was packed with a C18 reverse phase LiChroprep 40–63 μm (Lobar Merck, Darmstadt, Germany). After conditioning the stationary phase with methanol (4 × 250 mL, 0.1% formic acid) and distilled water (4 × 250 mL, 0.1% formic acid), 5 g of methanolic extract was dissolved in water and the mixture was deposited on the stationary phase. Elution of the sample was conducted using water (4 × 250 mL, 0.1% formic acid) in the first step and followed by methanol (4 × 250 mL, 0.1% formic acid) in the second step. This process yielded 1.4 g of processed methanolic extract. After condition optimisation at the analytical level, 50 mg of the extract were solubilized in 500 μL DMSO and injected using a Rheodyne® valve (1 mL loop). Semi-preparative HPLC-UV purification was performed on a Shimadzu system equipped with: LC20A module elution pumps, an SPD-20A UV/VIS detector, a 7725I Rheodyne® injection valve, and a FRC-10A fraction collector (Shimadzu, Kyoto, Japan). The HPLC system was controlled by the LabSolutions software. The HPLC conditions were selected as follows: Waters X-Bridge C18 column (250 × 19 mm i.d., 5 μm) equipped with a Waters C18 pre-column cartridge holder (10 × 19 mm i.d.); solvent system: ACN (2mM TEA) (B) and H_2_O (2mM TEA & 2mM ammonium acetate) (A). Optimized separation condition from the analytical was transferred to semi-preparative scale by a geometric gradient transfer software (Guillarme et al. 2008). The separation was conducted in gradient elution mode as follows: 5% B in 0-5 min, 12% B in 5-10 min, 30% B in 10-30 min, 60% B in 30-55 min, 100% B in 55-65 min. The column was reconditioned by equilibration with 5% of B in 15 min. Flow rate was equal to 17 mL/min and UV traces were recorded at 210 nm and 280 nm. The separation procedure yielded 0.3 mg of predicentrine and 3.4 mg of glaucine. Spectra for predicentrine (<u>CCMSLIB00005436122</u>) and glaucine (<u>CCMSLIB00005436123</u>) were deposited on GNPS servers.

#### 5.6.7 NMR analysis

The NMR spectra of each isolated compound was recorded on a Bruker BioSpin 600 MHz spectrometer (Avance Neo 600). Chemical shifts (δ) were recorded in parts per million in methanol‒ d4 with TMS as an internal standard. NMR data are available as Supplementary Material S1 and S2.

## Conflict of Interest

The authors declare that the research was conducted in the absence of any commercial or financial relationships that could be construed as a potential conflict of interest.

## Author Contributions

P-MA, AR, MDK and J-LW designed the study. JB wrote the python script for gnfinder output formatting. P-MA and AR wrote the scripts for dataset preparation, taxonomically informed scoring and results analysis. MDK, SO, MB and TS used the scripts for metabolite annotation and provided feedback. MB and SNE collected the *Glaucium* species. P-MA and AR performed the LCMS analysis on the *Glaucium* extracts. MB analyzed profiling data, isolated the compounds of *Glaucium* and established their structures. P-MA wrote the manuscript together with AR and J-LW. All authors discussed the results and commented on the manuscript.

## Funding

Jonathan Bisson gratefully acknowledge the support of this work by grant U41 AT008706 from NCCIH and ODS. The School of Pharmaceutical Sciences of the University of Geneva JLW is thankful to the Swiss National Science Foundation for the support in the acquisition of the NMR 600 MHz (SNF R’Equip grant 316030_164095).

## Acknowledgments

MDK acknowledges gratefully the support by Yamada Science Foundation and The Nagai Foundation Tokyo. The authors acknowledge Ali Bakiri for fruitful discussions on the optimization of the weights.

## Supplementary Material

See attached file

## Code and Data Availability Statement

Scripts and datasets generated and analyzed for this study can be found at the following OSF repository: https://osf.io/bvs6x/ (DOI 10.17605/OSF.IO/BVS6X)

## Notes

https://osf.io/bvs6x/

