## Supplementary Material for "Taxonomically informed scoring enhances confidence in natural products annotation"

The following Supplementary Material is available for this article:

### S1. Structural elucidation of predicentrine

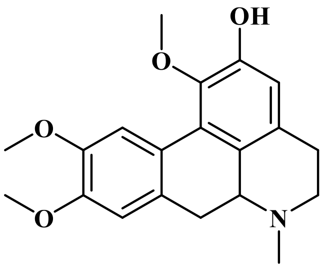

**Predicentrine:** ESI-HRMS (*m/z*): 342.1699 $[M + H]^+$; $^1$H NMR (**600 MHz,Methanol-$d_3$**) δ: 2.55 (1H, obsc, H-7b), 2.62 (1H,obsc , H-5b), 2.56 (3H, s, N–CH3), 2.70 (1H, dd, *J*=15.6, 4 Hz,H-4b), 3.12 (1H, obsc, H-6a), 3.14(1H, obsc, H-5a), 3.15 (1H, obsc, H-7a), 3.1 (1H, obsc, H-4a), 3.6 (3H, s, O–CH3), 3.88 (3H, s, O–CH3), 3.90 (3H, s, O–CH3), 6.6 (1H, s,H-3), 6.94 (1H, s,H-8), 8.03(1H, s,H-11), $^{13}$C NMR (**600 MHz,Methanol-$d_3$**), δ: 26.98 (C-4), 32.77(C-7), 41.62 (N-CH3), 52.06 (C-5), 54.22 ($C_9$-$OCH_3$), 54.45 ($C_{10}$-$OCH_3$), 58.31 ($C_1$-$OCH_3$), 148.78 (C-2), 61.87 (C-6a), 113.4(C-3), 110.56 (C-8), 110.98 (C-11), 123.67 (C-11a), 124.06 (C-6b), 125.58(C-11b), 127.74 (C-3a), 128.30 (C-7a), 142.50 (C-1), 147.2 (C-10), 147.50 (C-9). **(* Not assigned with certainty. "obsc.": The signal is obscured by overlapping peaks)**

**MSMS spectrum of predicentrine was deposited on the GNPS librairies (CCMSLIB00005436122).**

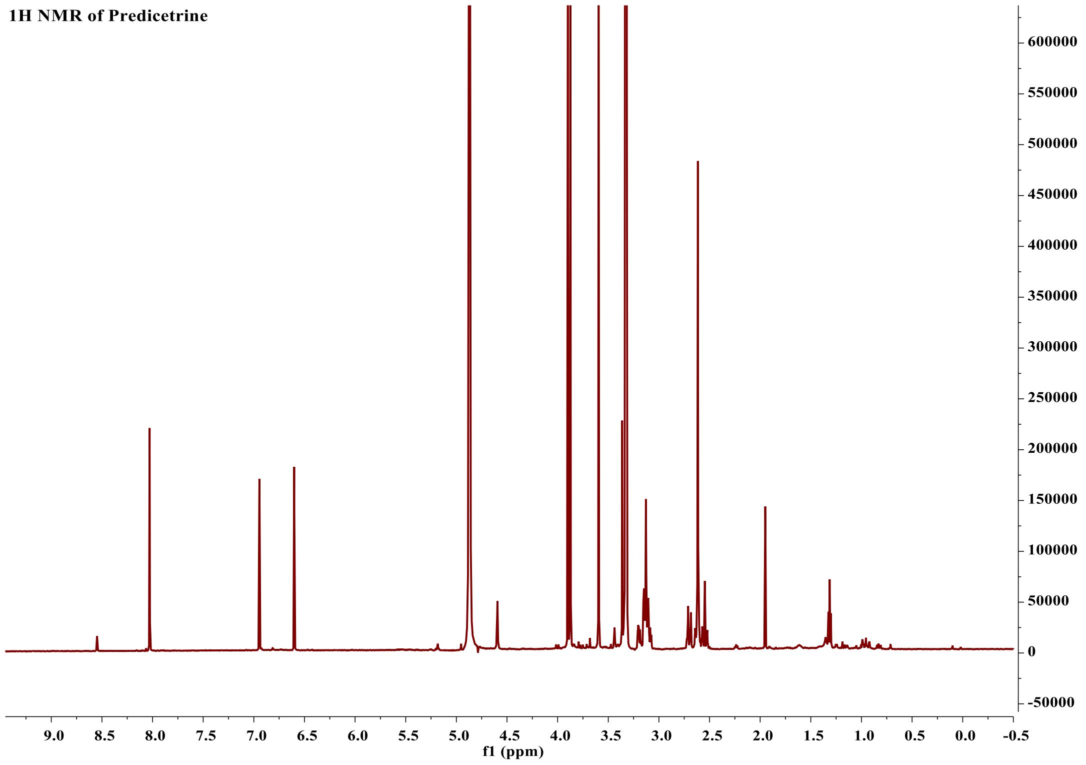

**Figure 1 1H NMR NMR spectrum of predicentrine**

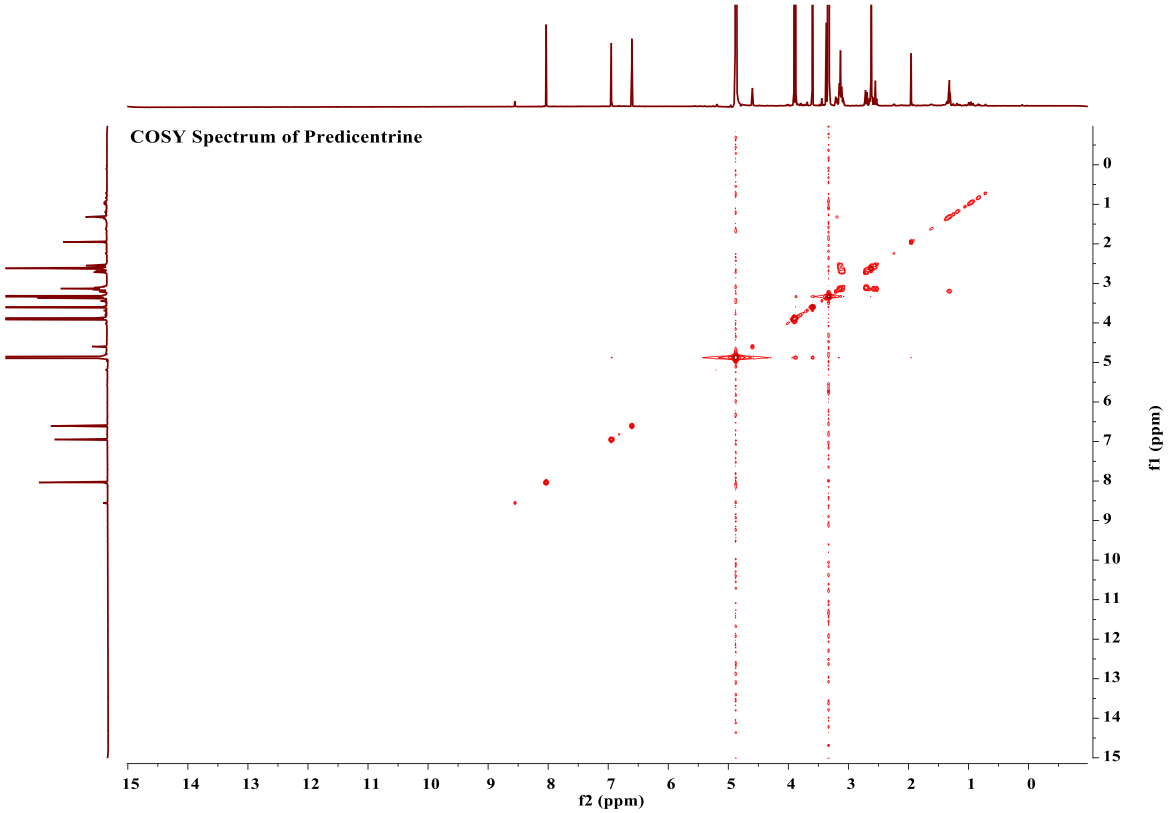

**Figure 2  COSY spectrum of predicentrine**

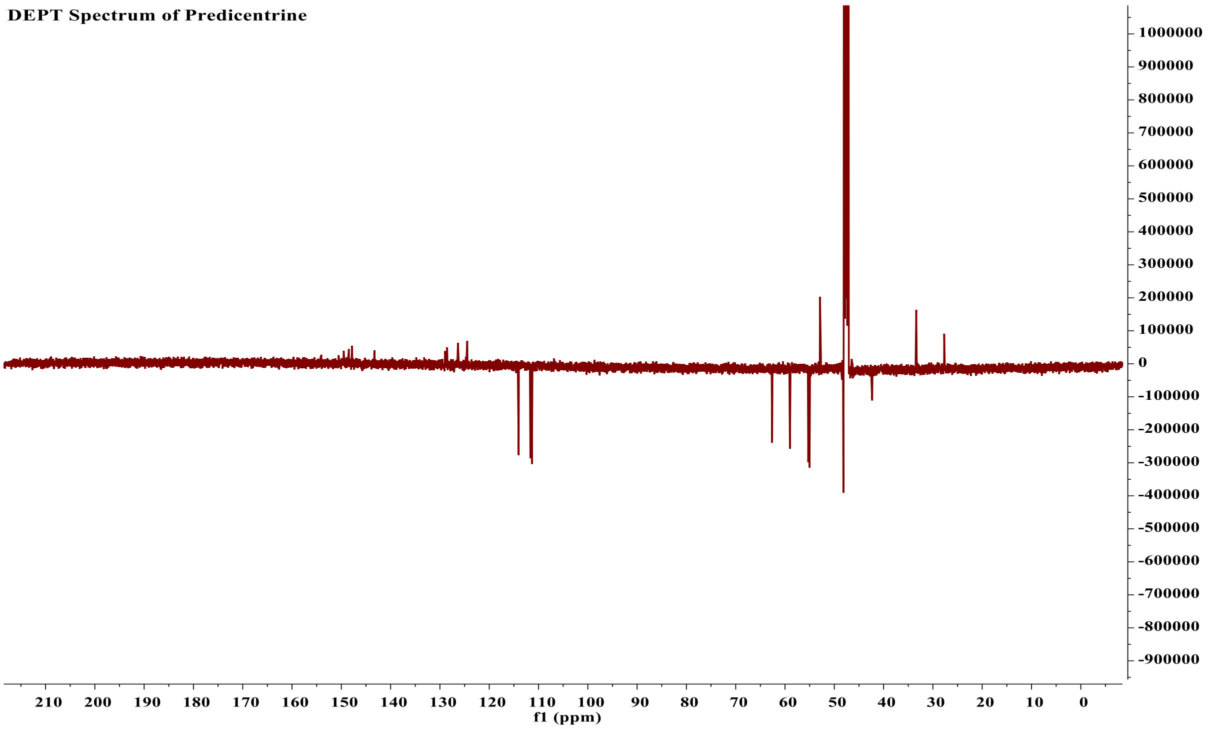

**Figure 3 DEPT spectrum of predicentrine**

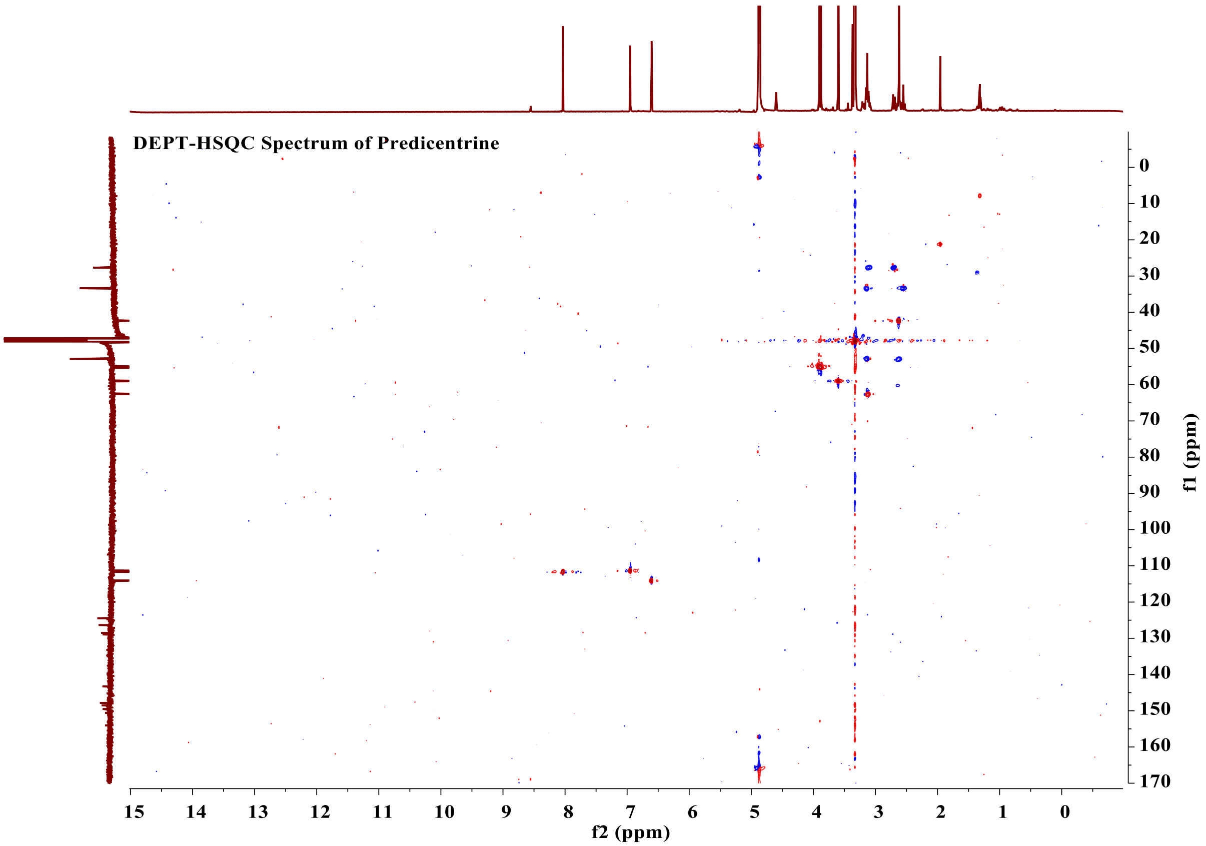

**Figure 4 DEPT-HSQC spectrum of predicentrine**

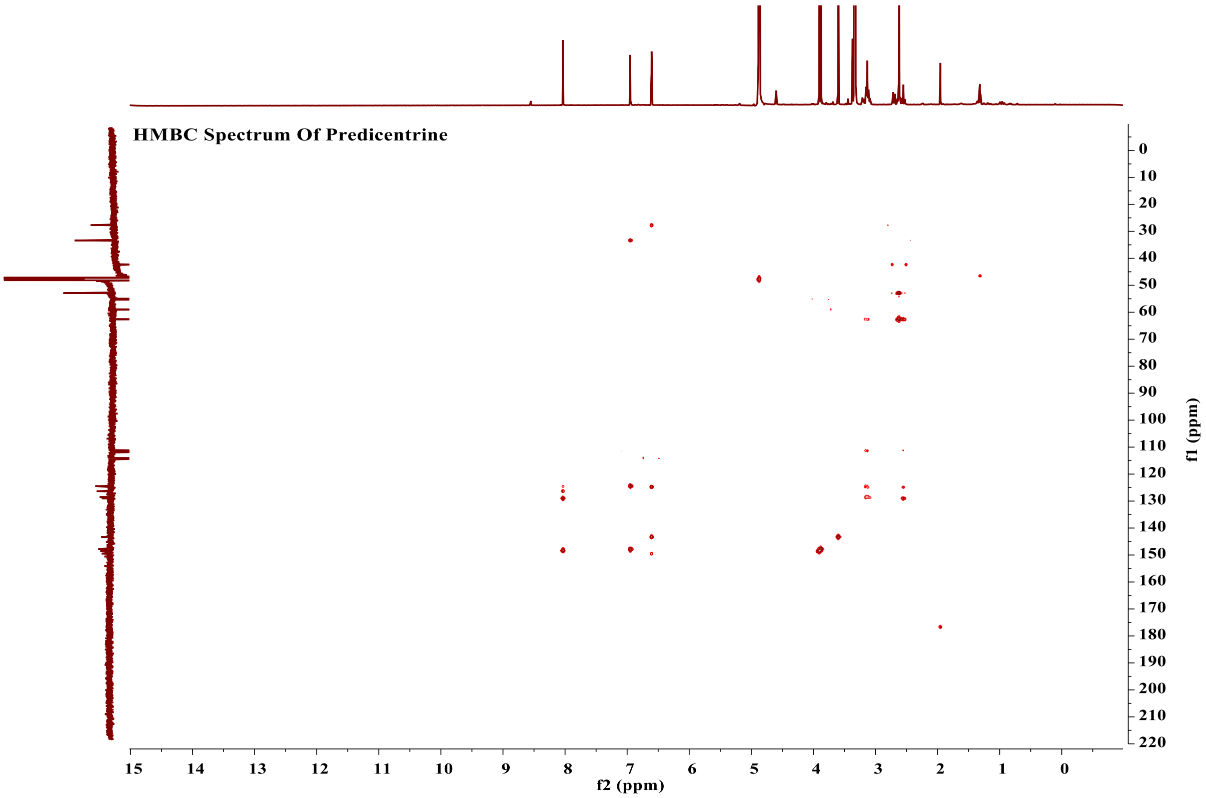

**Figure 5 HMBC spectrum of predicentrine**

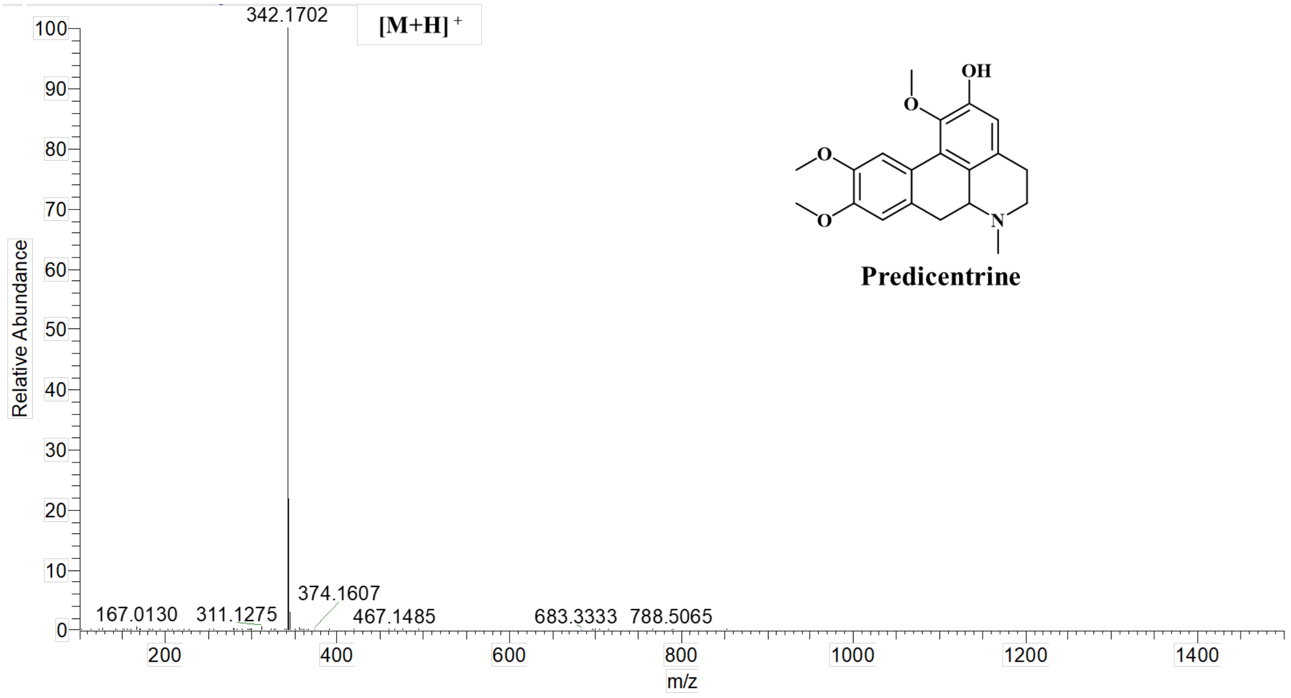

**Figure 6 HRMS spectrum of predicentrine**

### S2. Structural elucidation of glaucine

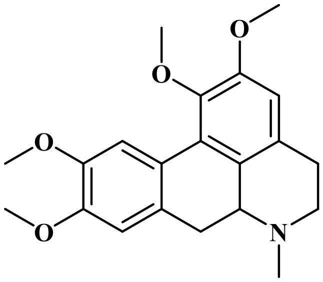

**Glaucine:** ESI-HRMS (*m/z*): 356.1859 [M + H]$^+$; $^1$H NMR (**600 MHz,Methanol-*d₃***) δ: 2.51 (1H, obsc, H-7b), 2.57 (1H, m, H-5b), 2.59 (3H, s, N–CH3), 2.75 (1H, dd, *J*=15.5, 3.7 Hz,H-4b), 3.07 (1H, dd, *J*=13.7, 3.9 Hz), 3.11(1H, obsc, H-5a), 3.14 (1H, obsc, H-7a), 3.15 (1H, obsc, H-4a), 3.64 (3H, s, O–CH3), 3.87 (3H, s, O–CH3), 3.88 (3H, s, O–CH3), 3.89 (3H, s, O–CH3), 6.74 (1H, s,H-3), 6.93 (1H, s,H-8), 8.01(1H, s,H-11), $^{13}$C NMR (**600 MHz,Methanol-$d_3$**), δ: 27.47 (C-4), 32.77(C-7), 41.79 (N-CH3), 52.17(C-5), 54.35 ($C_9$-$OCH_3$), 54.35 ($C_{10}$-$OCH_3$), 54.31 ($C_1$-$OCH_3$), 54.35 ($C_2$-$OCH_3$), 61.72 (C-6a), 109.55(C-3), 110.57 (C-8), 111.34 (C-11), 123.81 (C-11a), 151.68 (C-2), 125.36 (C-6b), 125.83(C-11b), 127.74 (C-3a), 128.55 (C-7a), 143.69(C-1), 146.83 (C-10), 147.81 (C-9). **(* Not assigned with certainty. "obsc.": The signal is obscured by overlapping peaks)**

**MSMS spectrum of glaucine was deposited on the GNPS librairies (CCMSLIB00005436123).**

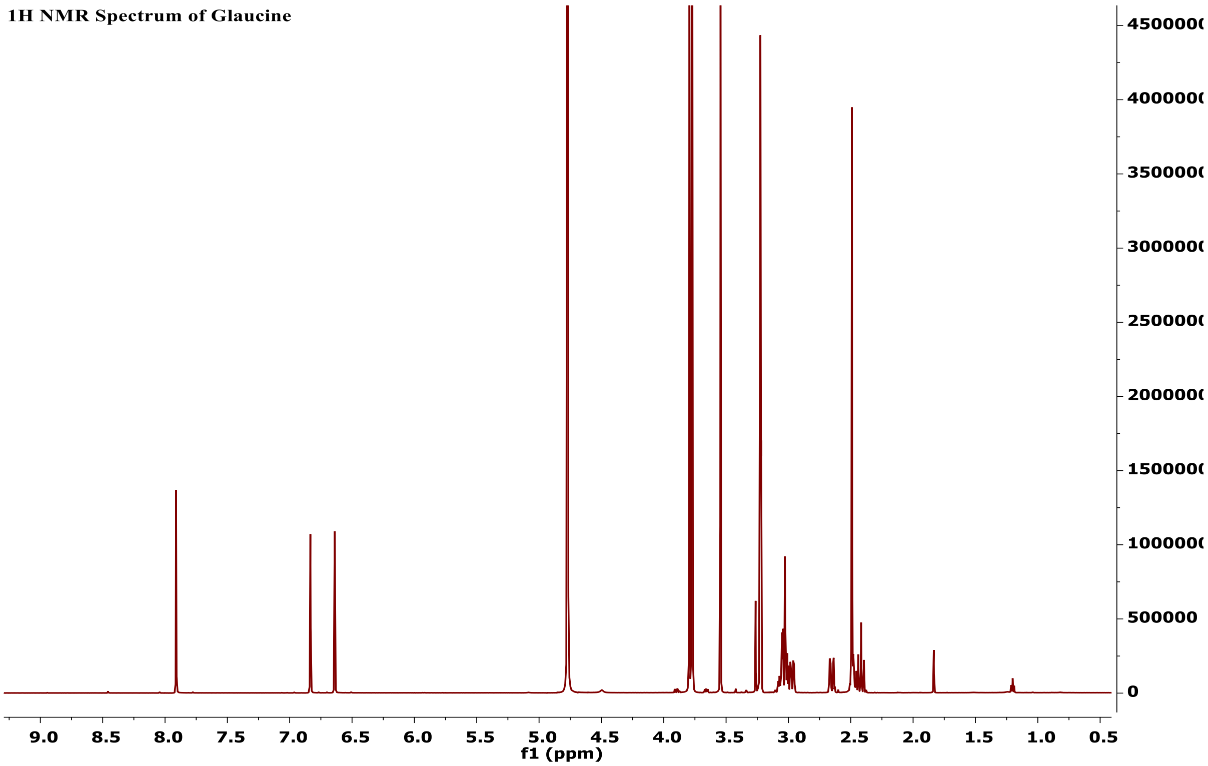

**Figure 7 1HNMR NMR spectrum of glaucine**

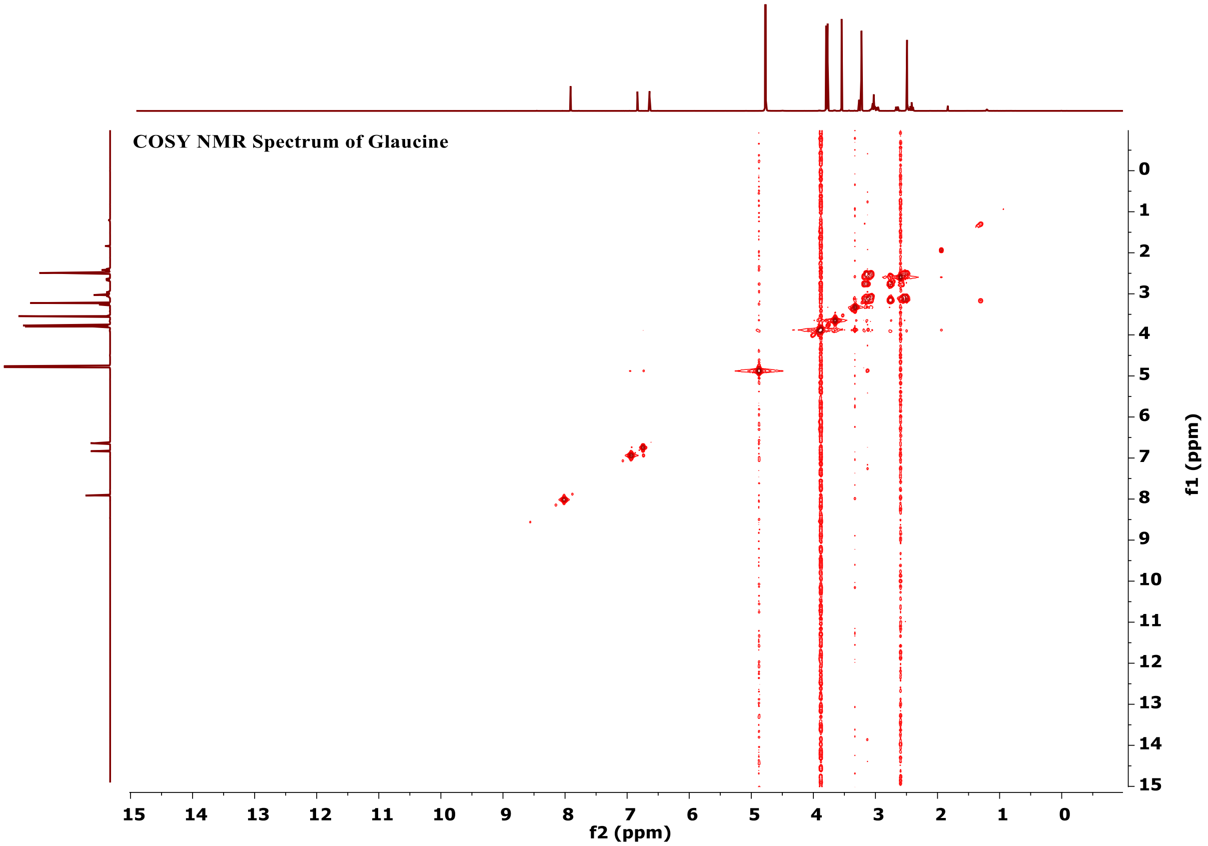

**Figure 8 COSY NMR spectrum of glaucine**

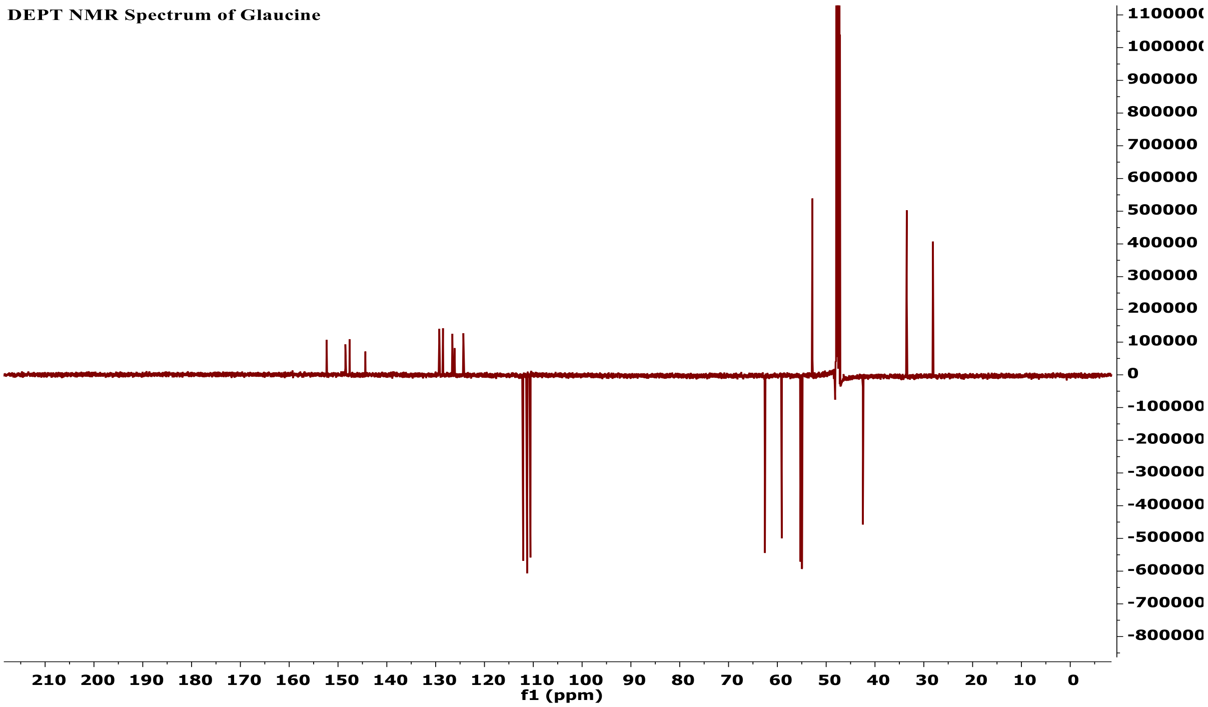

**Figure 9 DEPT NMR spectrum of glaucine**

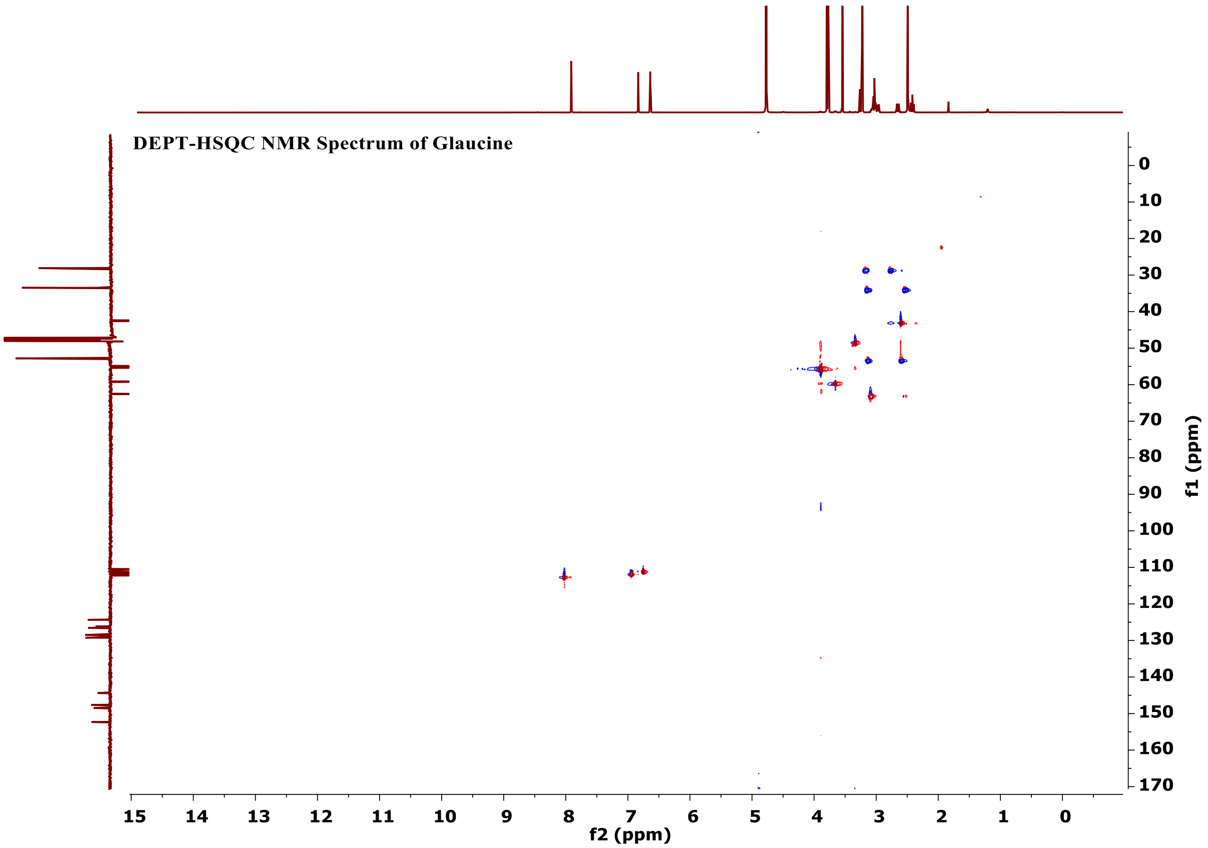

**Figure 10 DEPT-HSQC NMR spectrum of glaucine**

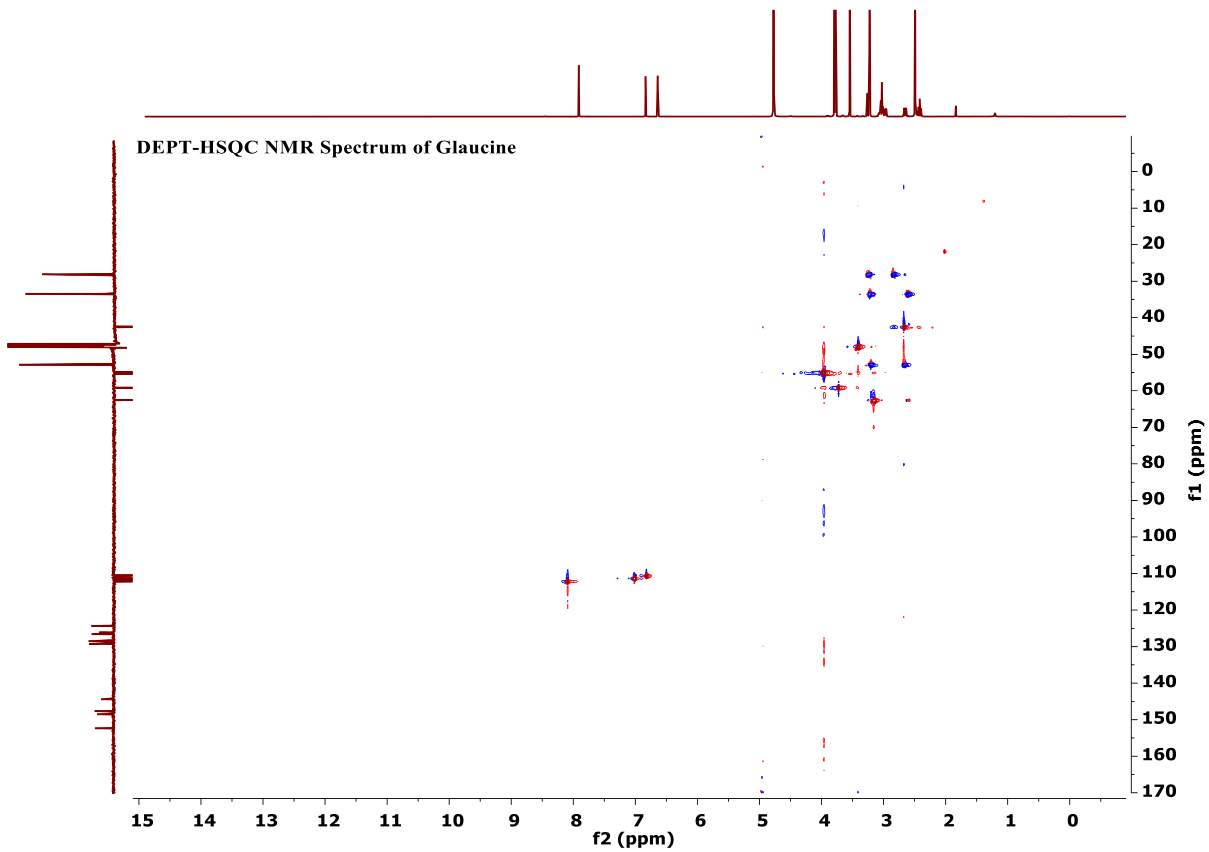

**Figure 11 HMBC NMR spectrum of glaucine**

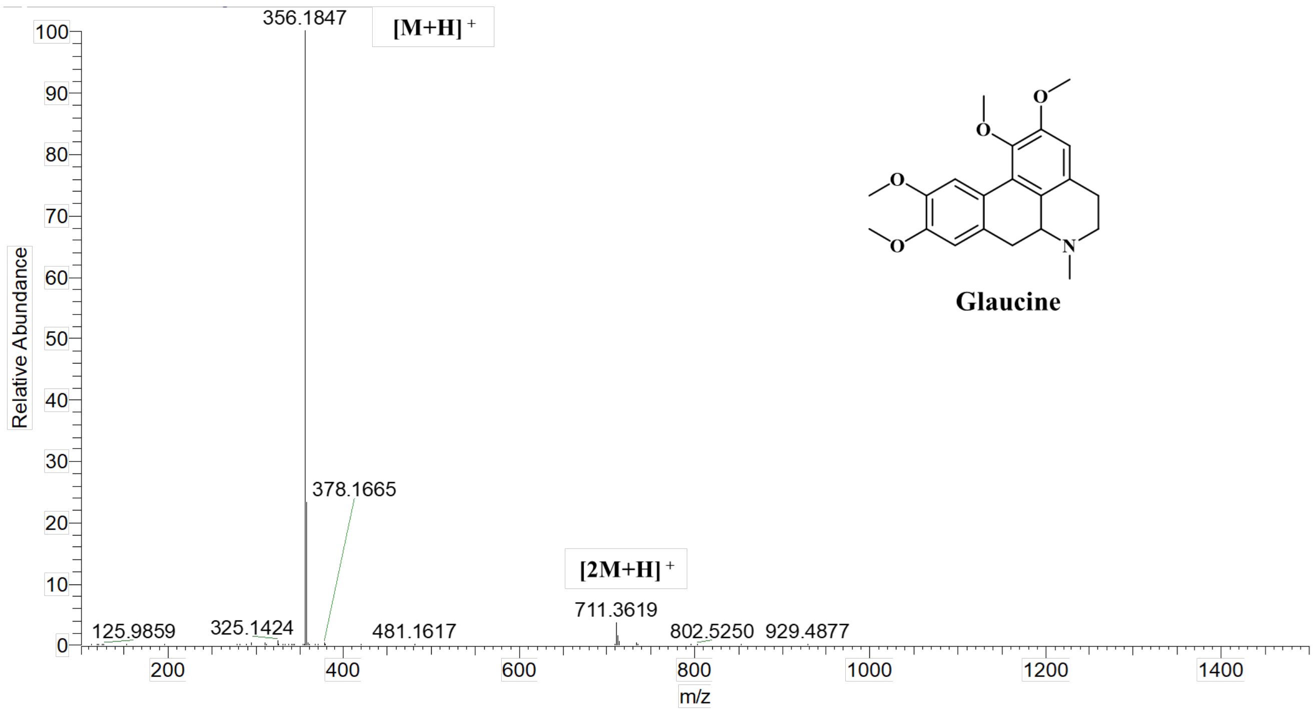

**Figure 12 HRMS spectrum of glaucine**

### S3. Entries corresponding to molecular formula $C_{16}H_{14}O_4$ in the benchmarking set

| Unique identifier | Short InChIKey | Family | Genus | Species | Identified |
|---|---|---|---|---|---|
| CCMSLIB00004702533 | ADRQFDIWPRFKSP | Fabaceae | *Bauhinia* | *Bauhinia guianensis* | Yes |
| CCMSLIB00004693536 | DEFIJGJJJKEYGS | Plagiochilaceae | *Plagiochila* | *Plagiochila adianthoides* | Yes |
| CCMSLIB00004694598 | IGWDEVSBEKYORK | Apiaceae | *Imperatoria* | *Imperatoria ostruthium* | Yes |
| CCMSLIB00000080466 | INBPQAJYHSJVRY | Betulaceae | *Alnus* | *Alnus sieboldiana* | Yes |
| CCMSLIB00000079496 | NSRJSISNDPOJOP | Fabaceae | *Dalbergia* | *Dalbergia nitidula* | Yes |
| CCMSLIB00000079436 | NYSZJNUIVUBQMM | Zingiberaceae | *Boesenbergia* | *Boesenbergia rotunda* | Yes |
| CCMSLIB00004697946 | OLOOJGVNMBJLLR | Apiaceae | *Imperatoria* | *Imperatoria ostruthium* | Yes |
| CCMSLIB00000077220 | ORJDDOBAOGKRJV | Zingiberaceae | *Boesenbergia* | *Boesenbergia rotunda* | No |
| CCMSLIB00004696878 | PACBGANPVNHGNP | Fabaceae | *Pisum* | *Pisum sativum* | Yes |
| CCMSLIB00004704345 | QQQCWVDPMPFUGF | Zingiberaceae | *Alpinia* | *Alpinia zerumbet* | Yes |

### S4. ROC curves (number of correct annotations vs. rank)

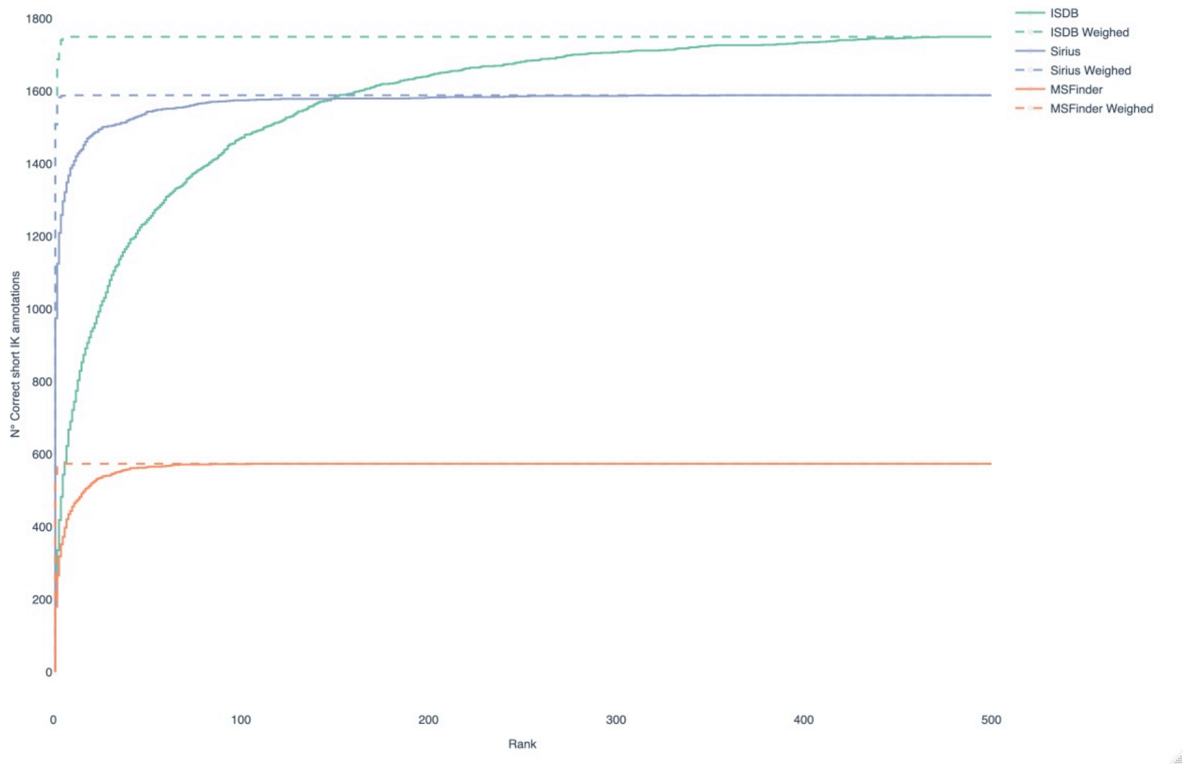

| Cluster ID | Structure | Short IK | Molecule Name | Family | Genus | Species | Family Weight | Genus Weight | Species Weight | Max Weight | Spectral Score | Normalized Spectral Score | Weighted Spectral Score | Rank Initial | Rank Weighted |
|---|---|---|---|---|---|---|---|---|---|---|---|---|---|---|---|
| **1771** | 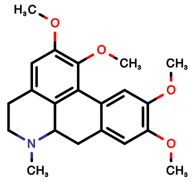 | **RUZIU YOSRD WYQF** | **1,2,9,10-Tetrahydroxyaporphine Tetra-Me ether (Glaucine)** | **Papaveraceae** | ***Glaucium*** | **NA** | **0.81** | **1.62** | **0** | **1.62** | **0.43** | **0.36** | **1.98** | **7** | **1** |
| **1771** | 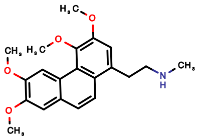 | QGNLU OSBJAG YFF | N-Methylsecoglaucine N-De-Me | Papaveraceae | *Corydalis* | *Corydalis yanhusuo* | **0.81** | 0 | **0** | 0.81 | 0.493 | 0.46 | 1.27 | 3 | 2 |
| **1771** | 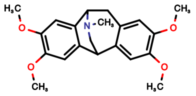 | KUHFD ZAEYV THRS | Thalisopavine Me ether | Papaveraceae | *Papaver* | *Papaver radicatum* | **0.81** | 0 | **0** | 0.81 | 0.455 | 0.4 | 1.21 | 4 | 3 |
| **1771** | 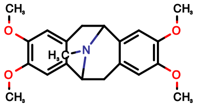 | QEOWC PFWLCI QSL | Argemonine | Papaveraceae | *Argemone* | *Argemone gracilenta* | **0.81** | 0 | **0** | 0.81 | 0.437 | 0.37 | 1.18 | 6 | 4 |
| **1771** | 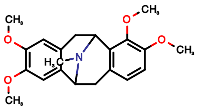 | PSKQB NMDFP YFNM | Platycerine Me ether | Papaveraceae | *Argemone* | *Argemone platyceras* | **0.81** | 0 | **0** | 0.81 | 0.428 | 0.35 | 1.17 | 8 | 5 |

**S5. Output of the taxonomically informed scoring annotation using ISDB-DNP for feature m/z 356.1860 at 1.83 min.** Glaucine, which is the correct annotation and was initially ranked at the 7th position, is now ranked at the first position.

### S6. Cluster related to predicentrine in *Glaucium* extract.

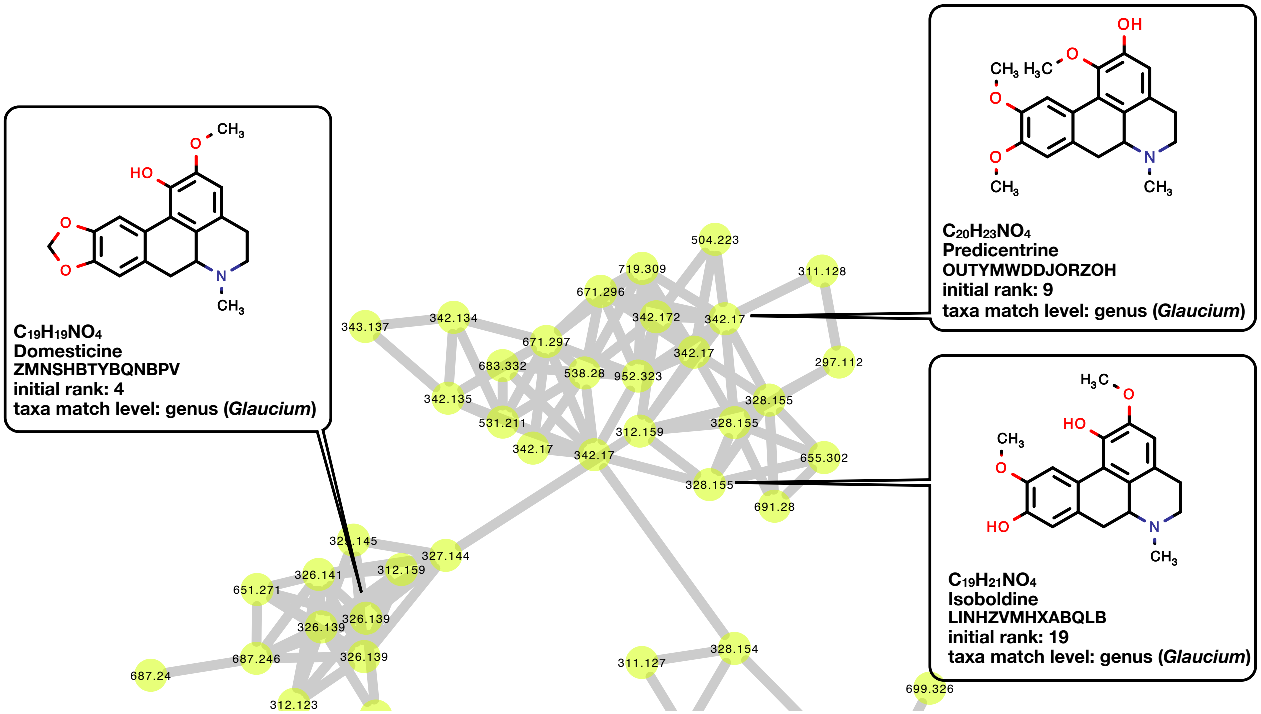

**Figure 13** Depiction of additional annotation (rank 1) returned by the taxonomically informed scoring applied on the *Glaucium* extract. Predicentrine was isolated, other annotations are putative. Full MN is available online: (https://gnps.ucsd.edu/ProteoSAFe/status.jsp?task=a475a78d9ae8484b904bcad7a16abd1f)
